# Avoiding tensional equilibrium in cells migrating on a matrix with cell-scale stiffness-heterogeneity

**DOI:** 10.1101/2020.10.15.342006

**Authors:** Hiroyuki Ebata, Satoru Kidoaki

## Abstract

Intracellular stresses affect various cell functions, including proliferation, differentiation and movement, which are dynamically modulated in migrating cells through continuous cell-shaping and remodeling of the cytoskeletal architecture induced by spatiotemporal interactions with extracellular matrix stiffness. When cells migrate on a matrix with cell-scale stiffness-heterogeneity, which is a common situation in living tissues, what intracellular stress dynamics (ISD) emerge? In this study, to explore this issue, finite element method-based traction force microscopy was applied to cells migrating on microelastically patterned gels. Two model systems of microelastically patterned gels (stiff/soft stripe and stiff triangular patterns) were designed to characterize the effects of a spatial constraint on cell-shaping and of the presence of different types of cues to induce competing cellular taxis (usual and reverse durotaxis) on the ISD, respectively. As the main result, the prolonged fluctuation of traction stress on a whole-cell scale was markedly enhanced on single cell-size triangular stiff patterns compared with homogeneous gels. Such ISD enhancement was found to be derived from the interplay between the nomadic migration of cells to regions with different degrees of stiffness and domain shape-dependent traction force dynamics, which should be an essential factor for keeping cells far from tensional equilibrium.

## Introduction

Changes in intracellular stress brought about by cell deformation and shaping have been shown to affect various cell functions. For example, cell contractility sensitively regulates DNA synthesis/cell proliferation^1, 2, 3, 4^, lineage-specific gene expression/cell differentiation^5, 6, 7, 8^, and the generation of cell-polarity/directed cell movement^9, 10, 11^. The strength and spatial distribution of intracellular stresses are determined by the interaction between mechanical properties of the extracellular matrix (ECM) and intrinsic cellular prestress which the cell actively generates inside itself as a result of the activities of adhesion machinery and cytoskeletal architecture ^12, 13, 14, 15^. In principle, this interaction is dynamic because of continuous cell-shaping and remodeling of the cytoskeletal architecture during cell movement. Since cells *in vivo* generally migrate in tissues with characteristic mechanical properties under a microscopic spatial distribution, intracellular stress dynamics (ISD) provide fingerprint-like information on the mechanical interactions between cells and ECM. Especially when the surrounding mechanical field is heterogeneous comparable to a cell-size scale^16, 17, 18, 19^, ISD in a single moving cell should exhibit characteristic long-term temporal fluctuations (i.e., range of hr) that are markedly different from those observed on a homogeneous mechanical field. Although this situation is rather common *in vivo*, the detailed characteristics of ISD on a matrix with cell-scale heterogeneity remain unclear.

To investigate this issue, there are two essential points to be addressed. One is the migratory behavior of a cell depending on the stiffness distribution on the matrix, which directly determines the ISD feature. The other is how to characterize long-term ISD on the heterogeneous field of stiffness. Concerning the first point, durotaxis, i.e., stiffness-gradient-dependent cellular taxis^20^, should be involved in determining ISD with the stiffness-heterogeneity. Conversely, the ISD feature can systematically be controlled through manipulation of durotaxis by designing the stiffness distribution on the microelastically patterned substrates^21, 22, 23, 24, 25, 26^. For example, in the case of striped-patterned gels with stiff and soft bands, usual soft-to-stiff durotaxis can be manipulated with the condition of elastic modulus and width of the bands^22^. An asymmetric elasticity gradient in the unit band of the stripe can induce unidirectional long-range ratchet-like rectification of durotaxis^27^. In addition, reverse stiff-to-soft durotaxis can be induced in a curved elasticity boundary that is convex toward the soft region with a curvature radius of less than 50 μm ^21^. Based on these cellular responses on durotaxis, microscopic stiffness-heterogeneity of ECM may simultaneously induce different types of durotaxis with a mixotactic manner^28^. To model this situation, the substrate that has both straight and curved stiffness boundary, such as a triangular stiff domain with dual tactic cues of usual durotaxis^20^ and reverse durotaxis^21^, should be an effective platform to characterize ISD on the matrix with stiffness heterogeneity (see Scheme 1).

**Scheme 1.**
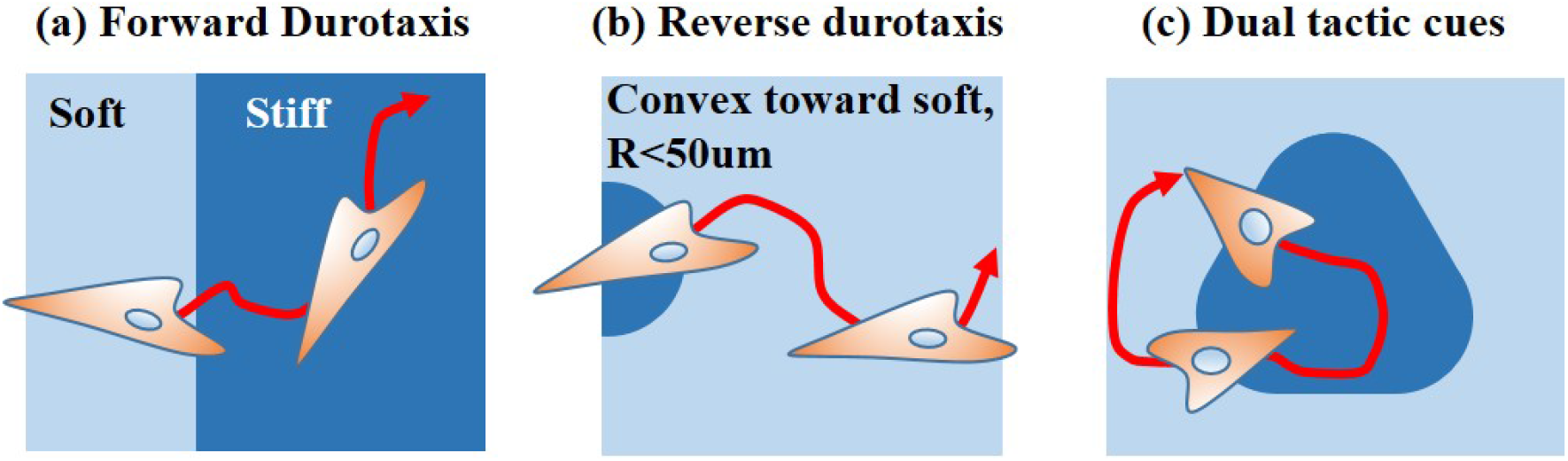
Manipulation of cell migrations by designing the matrix stiffness distribution. a) Usual linear stiffness boundary can induce forward durotaxis. b) Convex stiffness boundary toward the soft region with a radius of curvature smaller than 50 μm can induce reverse durotaxis^21^. c) Cyclic induction of forward and reverse durotaxis around the triangular stiff domain, the corner of which has a curved stiffness boundary to induce the reverse durotaxis.

As for the second point on the characterization of the ISD, traction force microscopy (TFM) is a powerful tool for determining the traction behavior at cell adhesion interfaces which reflect cellular contractility and intracellular stresses. Various types of TFM have been proposed for 2D and 3D analyses^29, 30, 31, 32^. When a cell adheres to a flat homogeneous gel surface, analytical solutions of linear elastic theory, the Boussinesq solution, are often used to reconstruct the traction force^29, 30^. However, for a matrix with cell-scale stiffness-heterogeneity, such as a microelastically patterned substrate, the elastic modulus sharply changes within the adhered area of a single cell, and an analytical solution for the equation of equilibrium cannot be obtained in general. Thus, for micromechanically-heterogeneous systems, TFM based on a numerical calculation such as with the finite element method (FEM) must be applied ^31, 33^. In addition, the time-scale for analysis is essential for investigating the effect of cell-scale stiffness-heterogeneity on ISD. In particular, the long-period fluctuation of traction force in an entire cell is important since it can cause field size-dependent periodic perturbation of the tensional steady-state of the cells. To analyze this phenomenon, TFM should be performed for more than several tens of hours.

In this study, to reveal the feature of ISD affected by cell-scale stiffness-heterogeneity, we sought to determine the long-term and whole cell-scale dynamics of traction force for cells migrating on a heterogeneous matrix. By employing FEM-based TFM^31, 33^, we measured the long-term dynamics of traction force of mesenchymal stem cells (MSCs) cultured on microelastically patterned hydrogels^21, 34^. Several tens of microns-scaled patterns were applied with stiff/soft stripes and periodic triangular stiff domains. In contrast to previous typical TFM results on a single linear elasticity gradient with pillar arrays^35, 36^, the size and curvature of the stiff domain were found to crucially affect the magnitude of the traction stress. We clarified that cell-scale variability of stiffness in an adhesion area determines the magnitude of the traction force as well as the local stiffness of the substrate. Reflecting these behaviors, the long-term computation of traction force dynamics revealed that cell-scale stiffness-heterogeneity strongly enhances the strength and frequency of fluctuation of the ISD in migrating MSCs in comparison to those on homogeneous gels, which was evidenced to keep the cells far from tensional equilibrium in the time scale of hours.

## Materials and methods

### Preparation of photocurable sol solution

Photocurable styrenated gelatin (StG) was used for photolithographic microelasticity patterning of the gel, which was prepared as described previously^21, 22, 34^. StG (30 wt %) and sulfonyl camphorquinone (3.0 wt % gelatin; Toronto Research Chemicals, ON, Canada) were dissolved in phosphate-buffered saline (PBS). The mixed solution was centrifuged (MX-301; TOMY, Tokyo, Japan) at 14,000 rpm (17,800 g) and 30 °C for 1h, and deposits were removed. The clear sol solution was aspirated for 1h at 45 °C to exclude dissolved oxygen. The sol solution was then conditioned for 10 min using an AR-100 deforming agitator (THINKY Corp., Tokyo, Japan).

### Photolithographic microelasticity patterning of gelatinous gel

Photolithographic microelasticity patterning gel with fluorescent beads was prepared as follows. First, a normal glass substrate was coated with poly(N-isopropylacrylamide) (PNIPAAm, Sigma Aldrich, St. Louis, MO) at 5 krpm for 60s. Next, we prepared diluted StG solution A; 10 wt % StG sol solution with 1.0 wt % sulfonyl camphorquinone and 0.1 wt % Tween 20. Sixty-five μl of the StG solution A was spin-coated on the NIPAAm-coated glass at 5 krpm for 20s. Next, we prepared diluted StG solution B; 10 wt % StG sol solution with 1.0 wt % sulfonyl camphorquinone, 0.1 wt % Tween 20, and fluorescent beads (Fluorospheres^®^ Carboxylate-Modified Microspheres, 0.2 μm, red fluorescence (580/605), Invitrogen). 65 μl of the StG solution B was spin-coated on the StG sol / NIPAAm-coated glass at 5 krpm for 20s. We added Tween 20 to improve wettability. Throughout the procedure, we warmed the glass and solutions at 45 °C to prevent dissolution of the NIPAAm layer. Next, 25 μl of the 30 wt % StG sol solution was spread between vinyl-glass and StG sol with beads / StG sol / NIPAAm-coated glass.

A soft base gel was prepared by irradiation of the entire sample with visible light for 120s – 210s (45 – 50 mW / cm^2^ at 488 nm; light source: MME-250; Moritex Saitama, Japan). Next, stiff regions were prepared by local irradiation of the base gel using a set irradiation pattern and a custom-built, mask-free, reduction-projection-type photolithographic system. Equilateral-triangular irradiated regions with side lengths *L* of 150 μm and 250 μm were projected on the gel using an EB-1770W liquid crystal display projector (SEIKO EPSON, Nagano, Japan). The center of the triangle was periodically placed on the honeycomb lattice. The distance *W* between the corners of the triangles was 120 μm (Fig. 1 (inset)). Finally, the gels were detached from the NIPAAm-coated normal glass substrate and washed thoroughly with PBS at 28 °C to remove the adsorbed PNIPAAm and Tween20. Stripe patterns with a unit length *L* of 100 and 600 μm (Fig. 1 (inset)) and homogeneous gels were also prepared. As we reported previously, the surface biochemical conditions were ensured to be the same for all the gel samples with different elasticities^24^.

**Figure 1.**
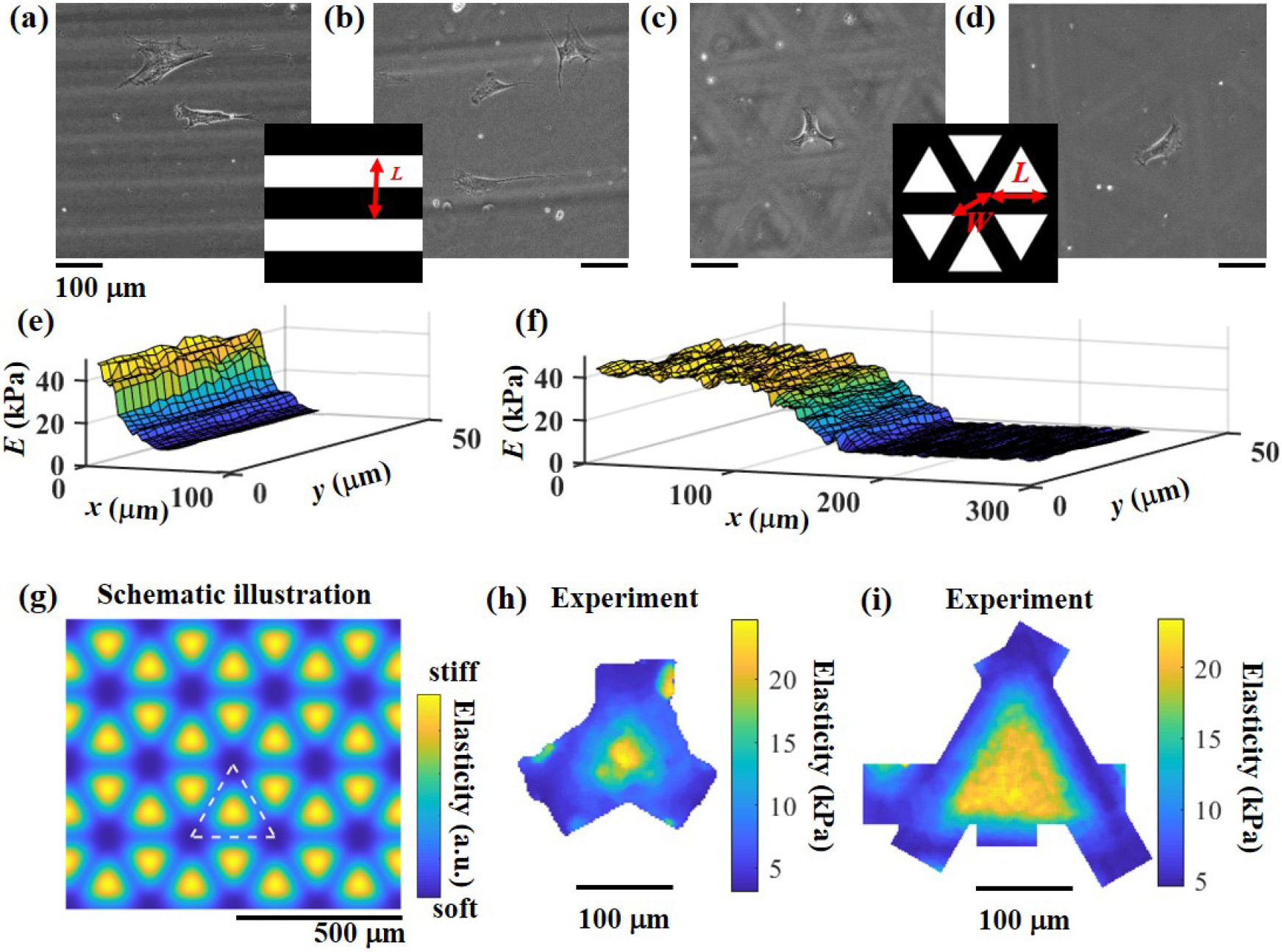
Striped- and triangular-patterned gels. **(a – d)** Phase-contrast microscopic images of Stripe-100 (a), Stripe-600 (b), Triangle-150 (c), and Triangle-250 (d). Scale bars denote 100 μm. (inset) Schematic illustration of the photomask. White indicates the irradiated region. Stripe: *L* is the unit width of a stripe. Triangle: *L* is the side length. *W* is the distance between the corners of the triangles, set at 120 μm. **(e, f)** Distribution of Young’s modulus *E* for Stripe-100 (e) and Stripe-600 (f). Distributions of a half unit-stripe are shown. **(g)** Schematic illustration of the spatial distribution of elasticity for triangular pattern. Blue and yellow colors indicate soft and stiff domains. The region surrounded by white dashed lines represents a unit triangle of the pattern. **(h, i)** Spatial distribution of Young’s modulus for Triangle-150 (h) and Triangle-250 (i).

### Measurement of the surface elasticity distribution around the elasticity boundary

The surface elasticity of the StG gel was determined by nano-indentation analysis using an atomic force microscope (JPK NanoWizard 4, JPK Instruments, Bruker Nano GmbH, Germany). A commercial silicone-nitride cantilever with a nominal spring constant of 0.03 - 0.09 N/m was used (qp-BioAC-CI CB3, Nanosensors). Young’s moduli of the surface were evaluated from force-indentation curves by nonlinear least-square fitting to the Hertz model in the case of a parabolic indenter (tip radius of curvature 30 nm; Poisson ratio μ: 0.5). To make a spatial distribution of elasticity in the triangular pattern gel, we measured about 100 tiles of a 40 × 40 μm elasticity map with a distance of 2 μm between each measured point. Next, we reconstructed the spatial distribution of elasticity from the tiles. We confirmed the linear-elasticity of StG gels through the measurement of the non-linearity of the elasticity and viscoelasticity (See SI, Fig. S1).

The topographic condition of the sample surface was also measured by using the atomic force microscope. Topography maps for microelastically patterned gels were shown in SI (Fig. S2). Although the soft regions were 15 – 25 μm higher than the stiff regions, the connection between the two regions was smooth enough to not disturb natural durotaxis. Compared with the cells on PDMS substrates^37^, we previously confirmed that cell migration was less affected by the topography of StG gels, and ~ 25 μm height difference did not induce any topotaxis (see SI, Figs. S2 and S3). In addition, from the topography map of homogeneous gels, we confirmed that the surface roughnesses were smaller than 80 nm irrespective of the stiffness (data not shown).

### Cell culture

Human bone marrow mesenchymal stem cells (hMSC, Lonza) were cultured in mesenchymal stem cell basal medium (MSCBM, Lonza) at 37°C in a humidified atmosphere containing 5% CO_2_.

### Time-lapse observation of cell movement

The migratory motion of cells on the microelastically patterned gel was monitored using an automated all-in-one microscope with a temperature- and humidity-controlled cell chamber (BZ-X700; Keyence Corporation, Osaka, Japan). Prior to the time-lapse observations, cells were seeded onto the gel surface at a density of 1 × 10^3^ cells / cm^2^ and cultured in DMEM containing 10% FBS for 18 – 22 hours under 5 % CO_2_. Phase-contrast images of cells and fluorescent images of beads were captured every 15 min for 24 h. We used a 20×(NA = 0.30) Plan Fluor objective lens. For each experiment, we collected images from 11 – 20 positions of 1 - 2 gel samples. For each position, we stacked the image of fluorescent beads for the *z*-direction. Next, we made an extended depth of field from the *z*-stacks using Image J^38^. To calculate the displacement field of the gels, we detached the cells after the time lapse. We replaced the medium with DMEM containing 0.3% Tween 20, and incubated the mixture for 30 min to kill the cells. Next, we washed the gel 3 times with DMEM. After incubation with DMEM for 30 min, we measured the reference image of the fluorescent beads.

### Analysis of cell-shaping dynamics

Movement trajectories and the shapes of the cells were determined and analyzed using MATLAB software. Based on edge-detection of a cell, we extracted the shape of the cell from phase-contrast images. The details of image-processing have been explained previously ^39^. We traced each cell and measured the time evolution of the trajectory and shape. If the cells collided with each other or if a cell was divided, we stopped the trace. When the cells separated again, we renumbered the cells and restarted the trace. Through the image analysis, cell trajectories *x*(*t*) were calculated.

### Traction force microscopy

To calculate the traction force of the cells, we used finite-element-method (FEM) based traction force microscopy^31, 33^. The displacement field of fluorescent beads was measured by comparing images with and without cells. The displacement of the beads was calculated using commercial PIV software (Flownizer 2D; Detect Corporation, Tokyo, Japan). In the calculation of FEM, a hexahedron was used for discretization (the size of the unit is about 8 × 8 × 8 μm). The nodes of a hexahedron at the substrate surface were chosen to be identical to the nodes of PIV. We assume that the patterned gel is a linear elastic body with spatial modulation according to Young’s modulus *E*(*x*):

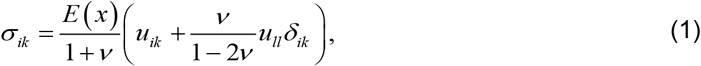

where *i*, *j*, and *k* are *x*, *y*, *z*. *σ_ik_* and *u_ik_* are the stress tensor and strain tensor, respectively. *v* is the Poisson ratio. In our calculation, we use *v*= 0.4. For homogeneous gels, *E*(*x*) is constant. For patterned gels, we assume

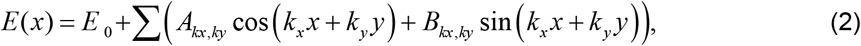

where *E_0_*, *A_kx,ky_*, and *B_kx,ky_* were fitted by the data from the AFM measurement. With FEM, the equilibrium equation for a linear elastic body, Eq. (1), is discretized as

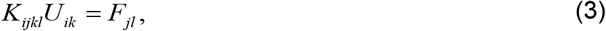

where *K_ijkl_* is a stiffness matrix, *U_ik_* is displacement of the *k* component at node *i*, and *F_jl_* is force in the *l* direction applied at node *j*. From Eq. (3), we calculate a response function *M_ijkl_* of the gel against the point force, where *M_ijkl_* is the *k* component of the displacement at node *i* when a unit force is applied in the *l* direction at node *j*. Since we do not observe the *z* direction of displacement, we only consider the *x* and *y* components in force reconstruction as in ordinary TFM in 2D. For simplicity, we introduce the following matrix and vectors.

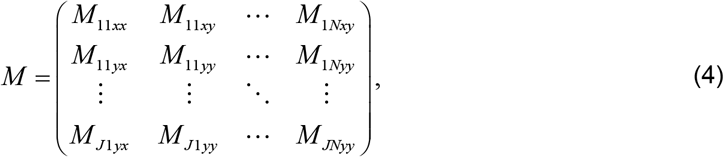

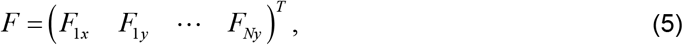

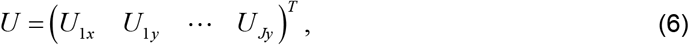

where ^T^ denotes the transposition. *N* is the number of the node where the force is reconstructed. *J* is the number of the node where the displacement is measured experimentally. In the case that the forces are distributed on surface nodes, displacement of the surface *U* can be expressed by using *M*:

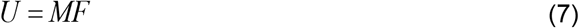

By solving the inverse problem of Eq. (7), we can reconstruct the distribution of the force. The flow chart of FEM-based TFM is shown in Fig. S4. Because of the ill posed nature of force reconstruction, some filtering schemes are required to reduce the noise. In this paper, *F* is reconstructed by minimizing function *S*, which is a combination of the least-square displacement error and a weighted norm of the forces:

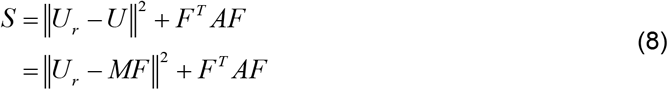

where *U_r_* is displacement of the substrate measured experimentally. The matrix *A* is a diagonal matrix containing the local penalty weights. Next, we calculate *F*, which gives the minimum of *S*, *dS*/*dF* = 0:

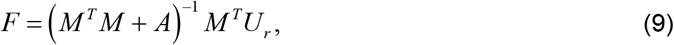

We then iteratively change *A* depending on the estimated force. We use a local penalty matrix that was proposed previously ^40^.

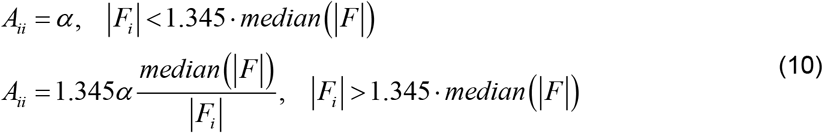

We repeatedly calculate Eqs. (9) and (10) until convergence is reached. When the regulation parameter *α* is sufficiently small compared to a typical value of *M^T^M*, the reconstructed traction field becomes very noisy. If *α* is too large, traction stress tends to be underestimated. Since *M^T^M* is inversely proportional to the square of the elasticity of the substrate, we vary *α* depending on the magnitude of *M^T^M*. We set regulation parameter *α* as

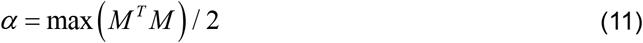

The validity and accuracy of FEM-based TFM are discussed in the SI (Figs. S5 and S6).

### Calculation of the force spot

To identify the force spot, we calculated the threshold of the traction stress. As reported previously ^41^, the cumulative distribution of the traction stress around the cell had exponential tails (Fig. S7):

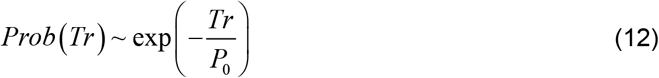

where the *P_0_* is the typical traction stress of the cell. *P_0_* does not depend on the segmentation of the cell shape. Typically, *P_0_* gave ~80% quantile of the traction stress under the cell. Then, we defined the domains where the traction stress was larger than *P_0_* as force spots. Thus, the top ~20% region of the traction stress was considered as force spots.

### Definitions of mean traction stress and mean elasticity

The mean traction stress *Tr_m_*(*t*) of a cell and mean elasticity *E_m_*(*t*) just below a cell are defined as

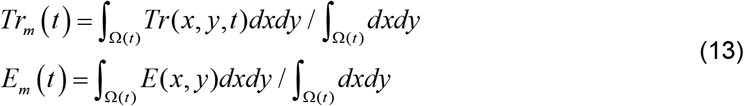

where *Tr*(*x*,*y,t*) and *E*(*x*,*y*) are the traction stress and elasticity at point (*x*,*y*). Ω(*t*) is the area of the cell that was calculated from the segmentation of the phase-contrast image.

### Definitions of magnitude of fluctuation of traction stress and other variables

Similar to the approach described in the literature^42^, we used the normalized standard deviation of *Tr_m_*(*t*) as the magnitude of fluctuation of traction stress. It is defined as the ratio of the standard deviation to the mean of the time series.

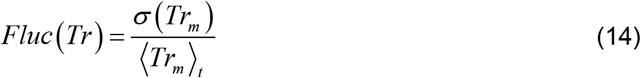

where σ(*Tr_m_*) and ⟨*Tr*_*m*_⟩_*t*_ are the standard deviation and time average of *Tr_m_*(*t*). The magnitude of the fluctuations of gel deformation, the cell aspect ratio, and cell area were calculated in a similar manner.

### Statistical analysis

For all figures, statistical analysis was performed by using the Mann–Whitney U test. P value between two groups of the data was calculated using Matlab software.

## Results

### Microelastically patterned substrates with cell-scale stiffness-heterogeneity

To investigate the effect of cell-scale stiffness-heterogeneity on cell migration and ISD, we prepared two types of microelastically patterned substrates by photolithography with photocurable styrenated gelatin (StG) (Fig. 1) ^21, 34^. One was a striped gel with alternating stiff and soft bands of different widths (targeted unit width *L*: 100 μm and 600 μm: Figs. 1a and b, designated Stripe-100 and Stripe-600, respectively). The other was a triangular-patterned gel with different side-lengths of a triangular stiff domain (targeted side length: 150 μm and 250 μm: Figs. 1c and d, designated Triangle-150 and Triangle-250, respectively). As controls, we also prepared homogenous soft and stiff gels, designated Plain-soft and Plain-stiff, respectively. To avoid the influence of the topography of the gel surface on natural cell movement, the focus condition of lithographic photoirradiation was optimized. The conditions and Young’s moduli of the stiff and soft domains are shown in Table 1.

**Table 1.**
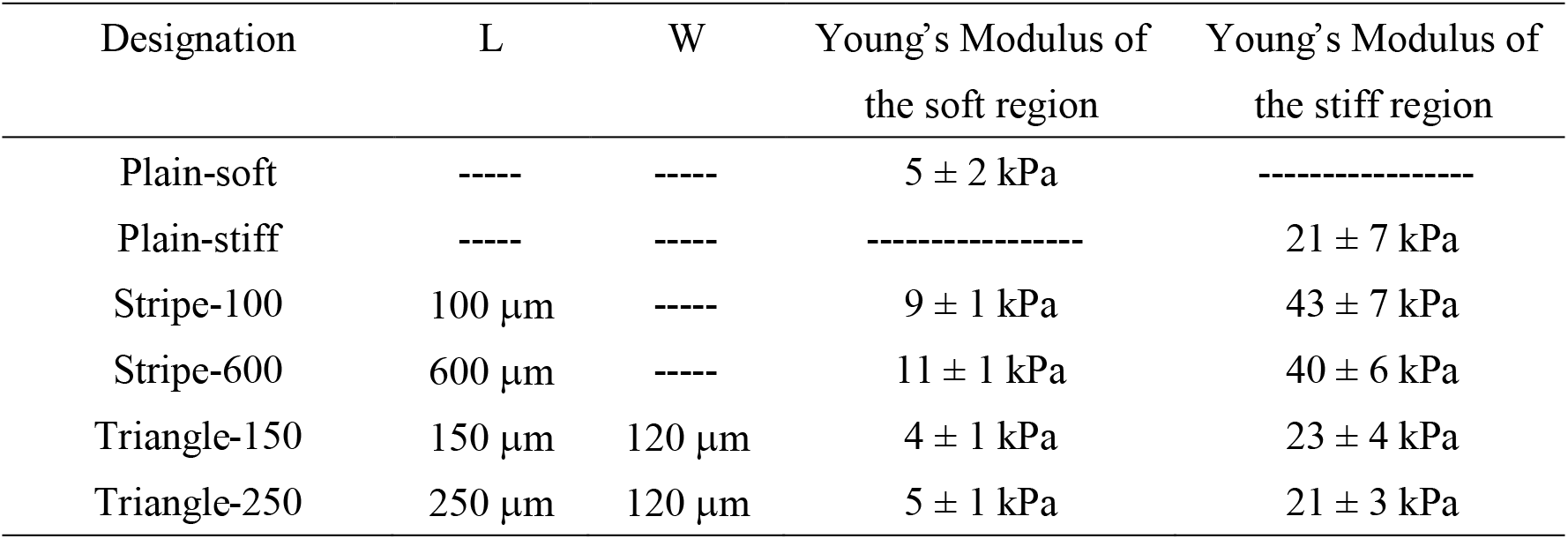
Size and Young’s moduli of the soft and stiff domains of microelastically patterned gels.

| Designation | L | W | Young's Modulus of<br>the soft region | Young's Modulus of<br>the stiff region |
| --- | --- | --- | --- | --- |
| Plain-soft | ---- | ---- | $5 \pm 2$ kPa | ----- |
| Plain-stiff | ---- | ---- | ----- | $21 \pm 7$ kPa |
| Stripe-100 | 100 $\mu\text{m}$ | ---- | $9 \pm 1$ kPa | $43 \pm 7$ kPa |
| Stripe-600 | 600 $\mu\text{m}$ | ---- | $11 \pm 1$ kPa | $40 \pm 6$ kPa |
| Triangle-150 | 150 $\mu\text{m}$ | 120 $\mu\text{m}$ | $4 \pm 1$ kPa | $23 \pm 4$ kPa |
| Triangle-250 | 250 $\mu\text{m}$ | 120 $\mu\text{m}$ | $5 \pm 1$ kPa | $21 \pm 3$ kPa |

For the striped gels, the unit widths of 100 μm and 600 μm were designed to model single cell-size and multicellular-size of MSCs, respectively. Young’s modulus was ca. 10 kPa for the soft regions and ca. 40 kPa for stiff regions over a transition region of ~50 μm in width (Figs. 1e and f). While a stiffness gradient of 30 kPa / 50 μm with a 10 kPa soft region was smaller than the threshold gradients of ~ 60 kPa / 50 μm to induce durotaxis on a single stiffness jump^43^, the narrow striped pattern has been previously clarified to enhance the induction of durotaxis depending on the cell-scale stiffness-heterogeneity^22^. Durotaxis is strongly induced in the 100 μm-wide stripes, but not in the 600 μm-wide stripes. To better understand the reason for these differential behaviors in durotaxis from the perspective of a traction force analysis, the present stiffness gradient was adopted.

For the triangular patterned gels, the side lengths of a stiff equilateral triangle *L* were designed to be 150 μm and 250 μm to model single cell-size and multiple cell-size, and the unit sizes of the as-prepared soft triangular base including stiff triangles were 254 μm and 354 μm, respectively. (Figs. 1c, d, and g). We changed *L* with a fixed width of soft regions *W* (see Fig. 1c and d inset). Young’s modulus varied from around 4 kPa to 20 kPa over a transition region of ca. 80 μm in width (Figs. 1h and i). The corner of a stiff triangle has a convex elasticity boundary toward the soft region with a curvature radius of less than 50 μm, where reverse durotaxis from stiff to soft regions can be induced, as we previously reported^21^.

### Stripe width-dependent modulation of traction force

To clarify ISD behavior on striped patterned gels with cell-scale heterogeneity^22^, traction forces were analyzed for the differential responses of durotaxis on different stripe widths with the same stiffness gradient. As shown in Fig. 2a, MSCs in the stiff regions of Stripe-100 exhibit usual durotaxis (blue). On the other hand, on Stripe-600, except in the transition region, they stayed almost equally in soft and stiff regions (red). On Stripe-600, cells tended to accumulate at the elasticity boundary, which has been explained by the model of durotactic migration^22^. Even though the strength of a stiffness gradient is set to be the same for these two conditions, only Stripe-100 induced durotaxis affected by the size of the stiff region. The mechanobiological basis for this differential induction of durotaxis was explored by using FEM-based TFM.

**Figure 2.**
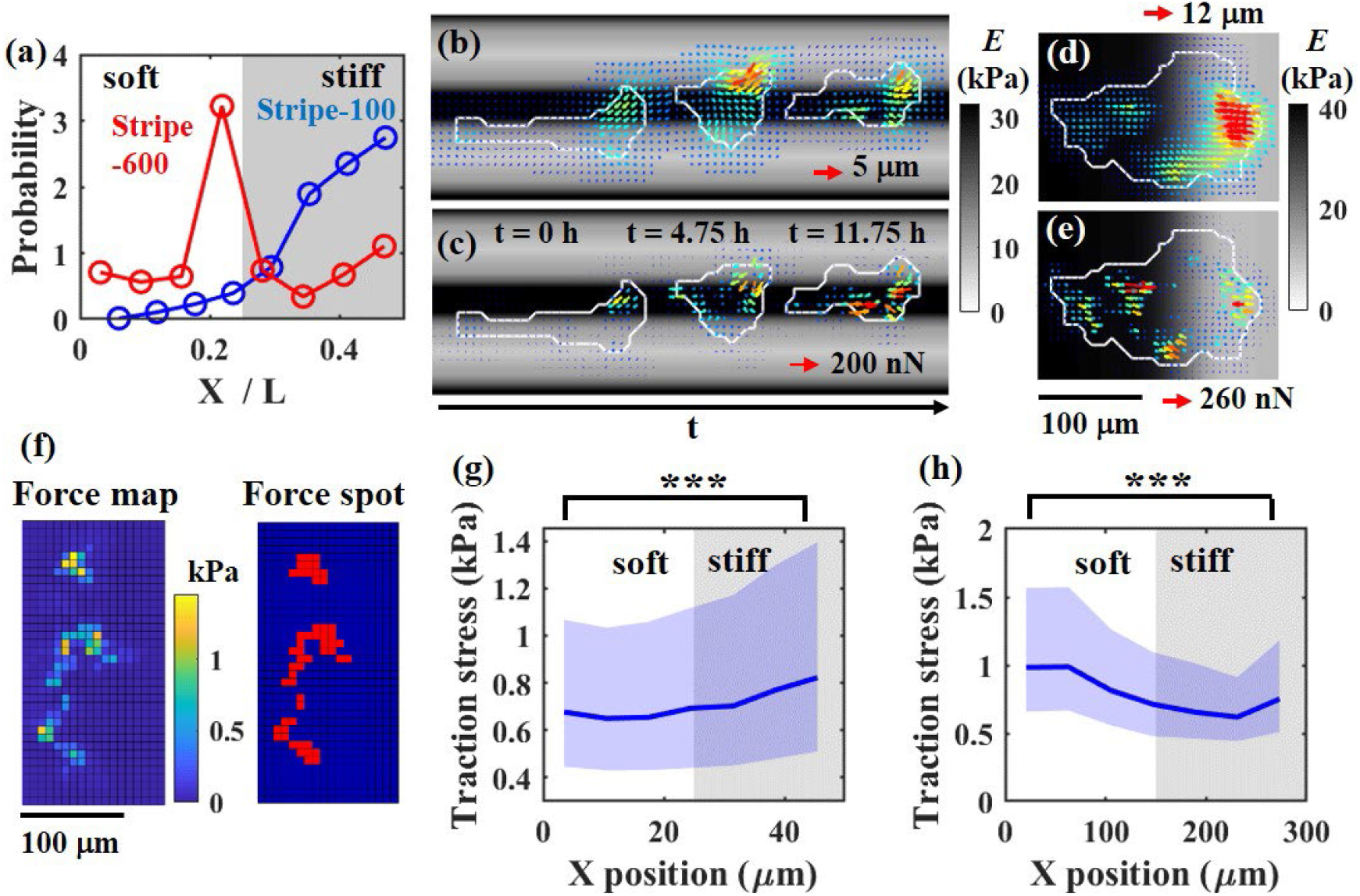
Dependence of traction force on the width of striped-patterned gels. **(a)** Probability distribution functions of the centroid of MSCs in a cycle of stripe. Horizontal axis denotes the position *x* normalized by the unit width *L* of the stripe. Blue: Stripe-100. Red: Stripe-600. **(b)** Deformation fields measured on Stripe-100. **(c)** Distribution of the traction forces estimated from Fig. 3 (b). **(d)** Deformation fields measured on Stripe-600. **(e)** Distribution of the traction forces estimated from Fig. 3 (d). (b - e) A vector field of different time points was superimposed on the figure. A longer vector represents greater deformation and force. As the color of the vector changes from blue to red, deformation and force increase. **(f)** An example of a color map of traction stress (left) and the corresponding force spots (right). **(g, h)** Magnitude of traction stress of the force spot as a function of the position *x* for Stripe-100 (g) and Stripe-600 (h). Solid lines indicate median values. Shaded regions cover 0.25 and 0.75 quantiles. The Mann–Whitney U test was used to calculate the P value. *** P < 0.001. n =32 for Stripe-100. n =25 for Stripe-600.

Figures 2b and c show a representative deformation field and reconstructed traction field generated by a MSC on Stripe-100. The background black/white shading represents the elasticity field of the gel. As the shading becomes dark, the gel becomes stiffer. Although the gel was highly deformed on soft regions (Fig. 2b), the magnitude of the traction force tended to be large on stiff regions (Fig. 2c). On the other hand, Figs. 2d and e show these fields on Stripe-600. Compared with the results for Stripe-100, large traction forces were exerted on both soft and stiff regions (Fig. 2e). These observations suggested that the striped pattern with a single-cell size could enhance the magnitude of traction force on a stiff domain.

This effect of stripe width on the traction force seemed to dominantly arise at the force spots where large traction stress was localized^41^ because the distribution of large focal adhesions was reported to overlap the domain of large traction stress. Therefore, we next analyzed the dependence of the force magnitude in the force spot on the spatial heterogeneity of stiffness. From the cumulative distribution of the traction stress of an entire cell, we calculated the threshold of traction stress to identify the force spots (Fig. 2f and Method). Typically, the top 20% regions of traction stress under the cell were considered as force spots. The left side of Fig. 2f shows an example of the spatial distribution of traction stress of a representative cell. Yellow domains on the right side of Fig. 2f represent the force spots calculated from the left side of Fig. 2f. Next, we calculated the magnitude of traction stress in force spots as a function of the position in the stripes (Figs. 2g and h). For Stripe-100, the traction stress of force spots became significantly large at the center of the stiff regions (right-hand edge of Fig. 2g). On the other hand, for Stripe-600, traction stress significantly increased in soft regions (left-hand edge of Fig. 2h). The smaller traction stress on stiff regions should not derive from the underestimation of the traction force during the force reconstruction process, because the traction stress of a force spot on the homogeneous stiff gel (0.48 kPa) became significantly larger than that on the homogeneous soft gel (0.38 kPa, see Fig. S8).

This analysis clarified that the traction forces in durotactic cells are sensitively modulated depending on the size of the stiff/soft regions even with the same stiffness gradient which does not originally induce durotaxis in a control linear elasticity boundary. A narrow stiff region enhanced the strength of traction stress enough to induce durotaxis, while a wide stiff region did not, which was found to be the cause of the emergence of differential responses of durotaxis for the same stiffness gradient.

### Traction forces around the triangular stiff domains

In the striped-patterned gels, one-way directed cell movement, i.e., soft-to-stiff durotaxis, was induced. To investigate ISD driven by dual tactic cues of usual durotaxis^20^ and reverse durotaxis^21^ as a model behavior of mixotactic cell migration in a living tissue, we analyzed traction forces on triangular patterned gels. First, we measured cell trajectories to characterize migration (Figs 3a and b). Here, the cell trajectory is defined as the path of the geometric center of the cell shape projected on an *x*-*y* plane, which was superimposed on the elasticity distribution estimated from AFM measurement; background black/white shading denotes stiff and soft regions, respectively. On Triangle-150, MSCs tended to go along the sides of stiff triangles and nomadically moved between soft and stiff regions around the corners of triangles, suggesting the emergence of dual tactic behaviors of usual and reverse durotaxis. Such frequent shuttling of MSCs was not observed in the striped pattern with a single-cell size (see Fig. 2a). On the other hand, on Triangle-250, MSCs tended to stay around the corner and the side of the triangle. As a result, MSCs stayed almost equally in soft and stiff regions on Triangle-150, while they stayed in stiff regions longer on Triangle-250 (Fig. S9). To confirm whether reverse durotaxis occurred, we calculated the probability distributions of cell position in the unit triangle of patterned gels, which was determined by the steady state of cell migration (Figs. 3c and d). On Triangle-150 (Fig. 3c), cells were distributed around the corners of triangles and migrated across the soft region between the corners of adjacent stiff triangles. On Triangle-250 (Fig. 3d), cells were strongly trapped near corner regions inside stiff domains.

**Figure 3.**
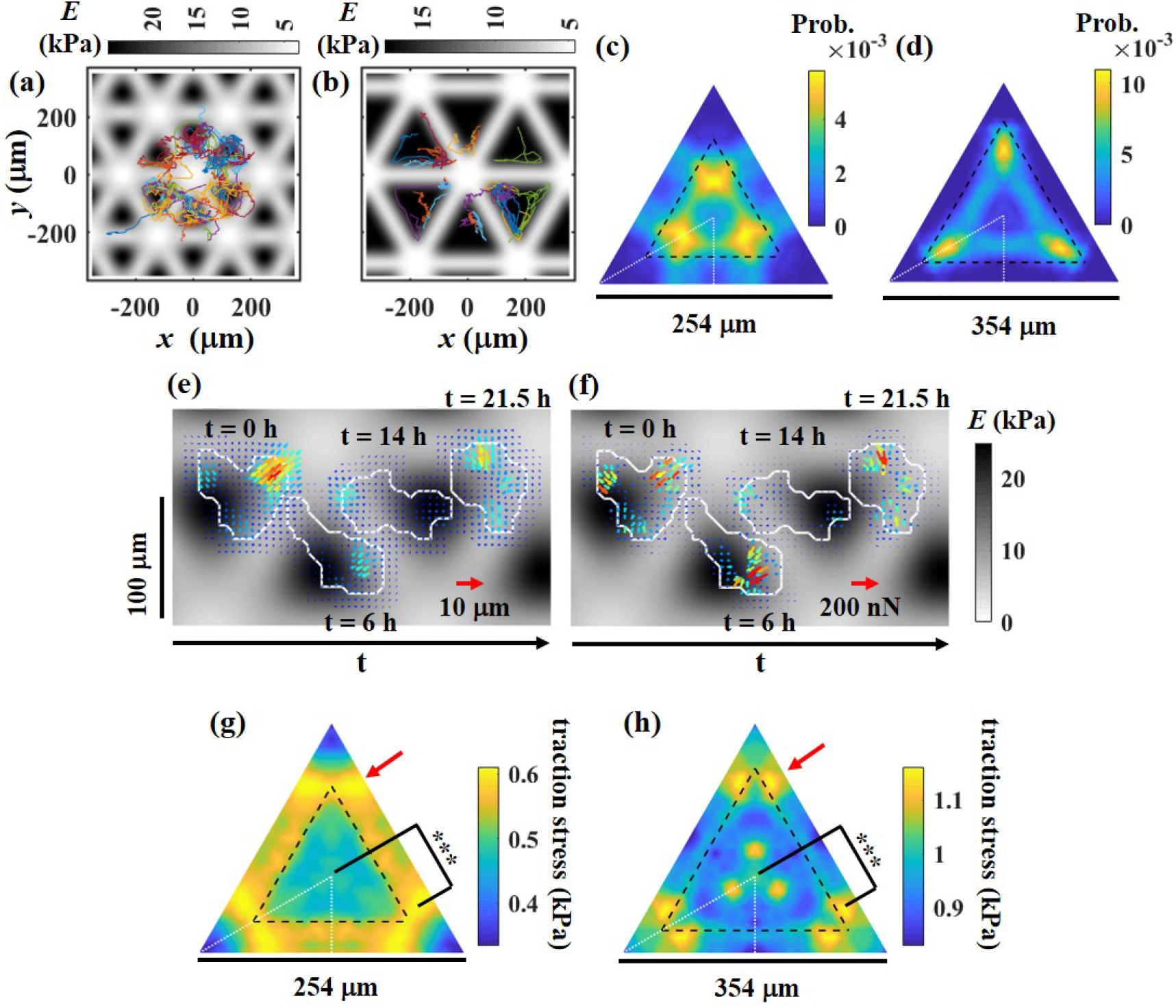
Dependence of the traction force on the convex shape of the stiff domain. **(a, b)** Trajectories of the centroid of the cells on Triangle-150 (a) and Triangle-250 (b). Gray-scaled background indicates the elasticity distribution of the gels. White and black indicate soft and stiff regions, respectively. Elasticity distribution was estimated from Figs. 1e and f. **(c, d)** Probability distribution of the cell position on unit triangular patterns of Triangle-150 (c) and Triangle-250 (d). Considering the spatial symmetry, the smallest stiff unit was surrounded by white dashed lines. We calculated the probability distribution on the smallest unit. Next, we replicated the smallest unit to construct the unit triangle. Thus, the probability distribution is highly symmetric. **(e)** Deformation fields measured on Triangle-150. **(f)** Distribution of the traction forces estimated from Fig. 4 (e). (e, f) A vector field of different time points was superimposed on a figure. A longer vector represents greater deformation and force. As the color of the vector changes from blue to red, deformation and force increase. **(g, h)** Color map of the traction stress of the force spot as a function of the position *x* and y. The Mann–Whitney U test was used to calculate the P value. *** P < 0.001. n =46 for Triangle-150. n =23 for Triangle-250.

What behaviors in traction forces emerge around triangular stiff regions? Figures 3e and f show the measured deformation field and reconstructed traction field generated by a MSC on Triangle-150. Interestingly, large traction forces of MSCs did not necessarily appear in the stiffest region, but rather appeared around the corners of triangles (Fig. 3f). The calculated spatial variation of the traction stress of the force spots (Figs. 3g and h) increased around the soft region outside of the corners of stiff domains for both small and large triangular patterns, indicating that the convex shape of the stiff domain crucially enhanced the magnitude of the traction stress in the soft region around the corners (Figs. 3g and h, red arrows). We clarified that MSCs are attracted to the soft outside of corners of stiff triangles by the effect of enhanced traction stress of the pseudopodia. In addition, Fig. 3h shows the increase in traction stress around the center of Triangle-250, suggesting the stronger induction of usual durotaxis toward stiff domains. This could cause the suppression of reverse durotaxis and nomadic motion on Triangle-250 (Fig. 3d). Thus, the results suggest that the size of the triangle modulated the balance of the competing cellular taxis.

### Temporal fluctuations of whole-cell traction stress affected by stiffness-heterogeneity

As clarified above, the spatial distribution and magnitude of traction forces in migrating cells are significantly modulated by the conditions of microscopic stiffness-heterogeneity, which, in principle, is a result of the characteristic dynamics in the remodeling of adhesion machinery and cytoskeleton on the matrix. Such dynamic remodeling of the intracellular architecture should cause modulations of ISD on long-term and whole-cell scales. To characterize the detailed features of the ISD, we analyzed temporal fluctuations of traction forces (Fig. 4).

**Figure 4.**
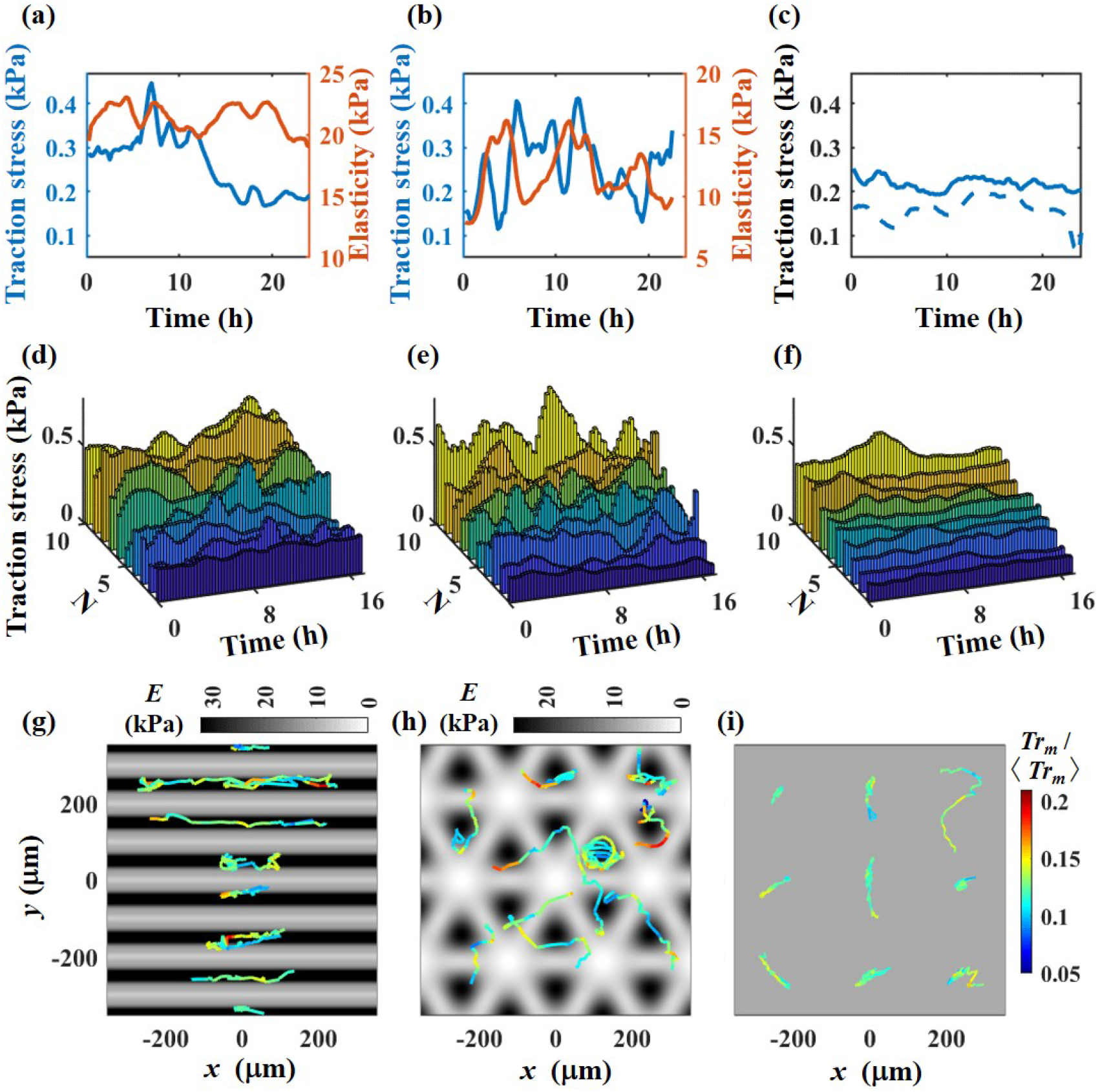
Time course of the traction stress on the patterned and homogeneous gels. **(a - c)** Example time course of the mean traction stress (blue) of a cell and mean elasticity (orange) under a cell on Stripe-100 (a), Triangle-150 (b), Plain-soft (c; dashed line), and Plain-stiff (c; solid line). **(d – f)** Time course of the mean traction stress of 10 cells on Stripe-100 (d), Triangle-150 (e), and Plain-stiff (f). Different colors indicate different cells. *N* denotes the cell number. (g – i) The magnitude of the traction stress was projected on the trajectories of the cells. The colors of the trajectory represent normalized mean traction stress. (g) Stripe-100. (h) Triangle-150. (i) Plain-stiff.

For patterned gels, the time series of mean traction stress was found to largely fluctuate during cell movement between soft and stiff regions (Figs. 4a and b, blue lines) compared to the control (Fig. 4c, blue lines). This tendency was confirmed for 10 cells in each condition (Figs. 4d-f), suggesting that fluctuation of the mean traction stress was greatest on triangular gels and least on the homogeneous gel. In the case of striped gel (Fig. 4d), they seemed to be intermediate. To simultaneously show the fluctuation of traction stress and positional information of the cells, we projected the time course of mean traction stress on the trajectories of the cells. In Figs. 4g-i, the colors of the trajectories represent the magnitude of normalized mean traction stress. The fluctuation of traction stress was moderate on Stripe-100, where the cells were strongly trapped in stiff regions (Fig. 4g). On the other hand, traction stress on Triangle-150 largely fluctuated when the cells passed through the side and corner of the stiff triangles (Fig. 4h). Compared with the cells on the patterned gels, fluctuation of traction stress was small on Plain-stiff gel (Fig. 4i).

To characterize the typical time-scale of the observed fluctuations of traction stress, we calculated the population-averaged autocorrelation function of the time series of traction stress (Fig. 5a). The autocorrelation functions exponentially decayed to zero within 2 – 4 hours without a clear peak, indicating that the fluctuation of the traction force was rather noisy. Through fitting of the autocorrelation function by an exponential exp(−*κt*), the correlation time *T_f_* = ln2/*κ* for each cell on a triangular patterned gel was found to be about 50 % smaller than that on homogeneous gels (Fig. 5 b). This result quantitatively confirmed that the temporal fluctuation of traction stress is strongly enhanced on triangular patterned gel.

**Figure 5.**
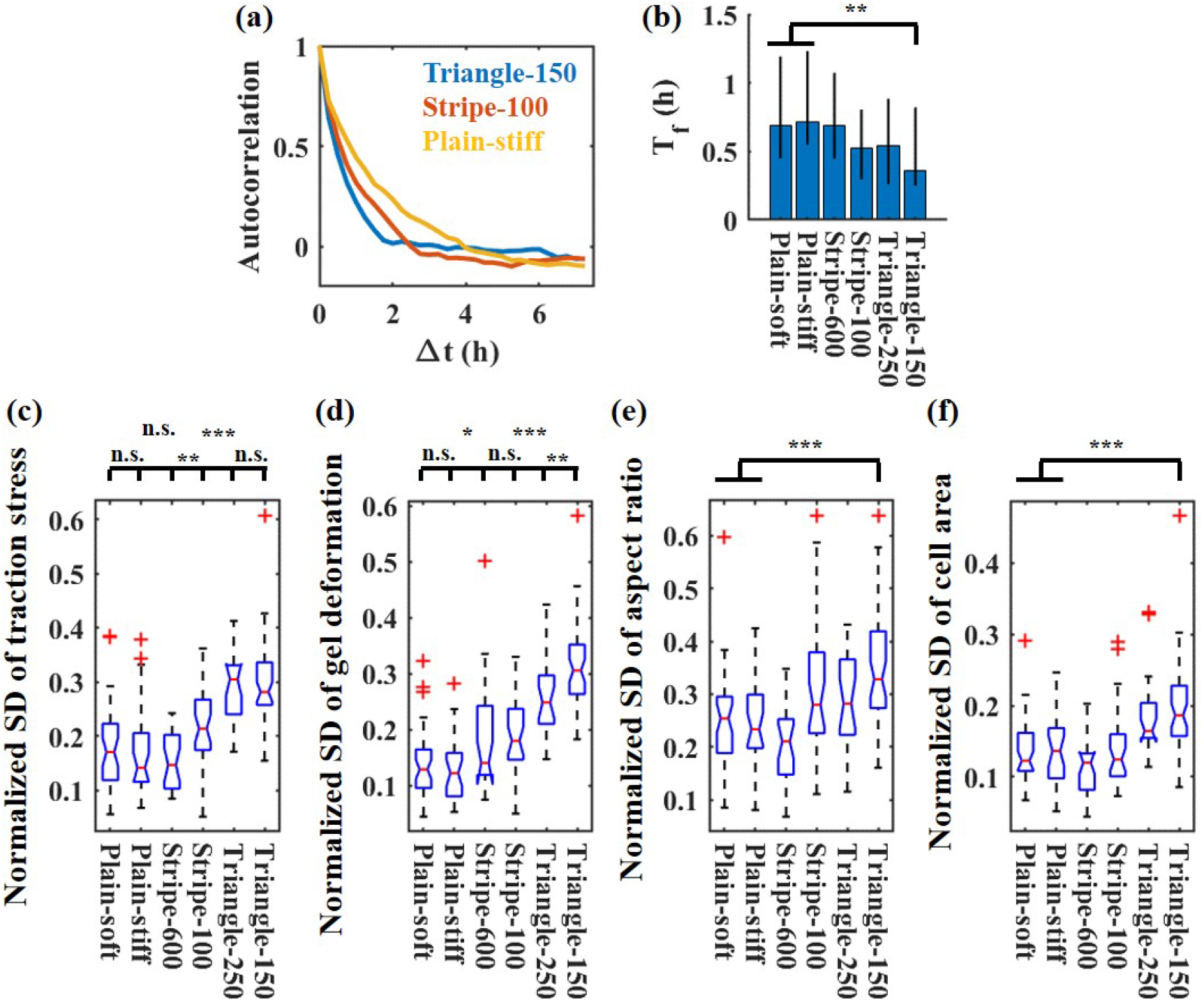
Quantification of fluctuation of mean traction stress and cell shapes. **(a)** Autocorrelation function of the time course of mean traction stress. Red: Stripe-100. Blue: Triangule-150. Orange: Plain-stiff. **(b)** Bar graph of correlation time of mean traction stress. Bar denotes a median value. Error bar connects the 0.25 and 0.75 quantiles. **(c)** Normalized standard deviation (SD) of the traction stress calculated from the time course of mean traction stress of a cell. Standard deviation of the traction stress was divided by the time-averaged traction stress. **(d - f)** Normalized SD of the mean deformation field of the gel just under a cell (d), aspect ratio (e), and cell area (f). Standard deviations of the parameters were divided by the time-averaged values. The Mann–Whitney U test was used to calculate the P value. * P < 0.05. ** P < 0.01. *** P < 0.001. Plain-soft: n = 39. Plain-stiff: n=28. Stripe-600: n = 25. Stripe-100: n = 32. Triangle-250: n = 23. Triangle-150: n = 46.

Regarding the magnitude of the temporal fluctuation of traction stress, the standard deviation of the time series of traction stress divided by the average traction stress of each cell (see Method) was about two times larger on triangular patterned gels than on homogeneous gels (Fig. 5 c). The fluctuation on Stripe-100 was intermediate between those on triangular and homogeneous gels. No significant difference was noted between Stripe-600 and homogeneous gel. Similar to the fluctuation of the traction stress, the magnitude of the deformation strongly fluctuated on the triangular pattern gels (Fig. 5 d). The large fluctuation of the traction stress should couple with the fluctuation of cell-shaping in correlation with the cell polarization and adhesion. The measured fluctuation of both the cell aspect ratio and area was largest on the triangular patterned gel with small stiff regions (Figs. 5e and f, respectively). In addition, we found that the cells remembered the nomadic migration on the triangular patterned gels for a few hours, and the cells kept large fluctuation of traction stress even when they temporarily stayed in a region with constant stiffness (see SI, Fig. S10).

As revealed above, on the matrix with cell-scale stiffness-heterogeneity, traction stress dynamics exhibit characteristic fluctuations depending on the microscopic patterns of stiffness, and especially oscillate between low and high values when the cell body repeatedly intersects the micro-pattern every few hours, as demonstrated around the triangular stiff domains. In conclusion, the matrix with dual cues for competing taxis on cell-scale stiffness-heterogeneity was confirmed to induce extraordinarily large and frequent fluctuation of ISD together with enhanced morphological fluctuations.

## Discussion

In this study, we focused on the features of intracellular stress dynamics (ISD) in cells migrating on a matrix with cell-scale stiffness-heterogeneity. To address this issue, two model systems of microelastically patterned gels were used: stiff/soft stripe and stiff triangular patterns. These stripe and triangular patterns were designed to characterize the effects of a spatial constraint for cell-shaping and of different coexistent cues to induce competing cellular taxis (usual and reverse durotaxis) on the ISD, respectively. One of the most striking ISD responses in migrating cells was that the strength of long-term fluctuation of traction force on a whole-cell scale was markedly enhanced in the latter system of triangular patterns compared with a stiffness-homogeneous matrix, which made the cells far from tensional equilibrium. Based on this finding, we discuss the following issues: (1) why is the fluctuation of traction forces enhanced on the triangular patterned system?, and (2) what kind of functional effect would migrating cells receive by remaining far from tensional equilibrium?

First, why was the fluctuation of traction stress amplified, especially on triangular patterns? In principle, large variations in the development and decline of traction forces in an entire cell should involve a sufficient residence time of cells on the stiff and soft regions for the development and dissipation of forces. In addition, the residence times should be balanced between these regions through nomadic cell migration to attain the up-and-down cycles of traction force dynamics (see Fig. S9): i.e., nomadic migration on a matrix with stiffness-heterogeneity is the essential cause of the amplification of the fluctuation of traction force dynamics. A matrix with homogeneous stiffness cannot induce cyclic force development and dissipation. Even though the matrix has a heterogeneous stiffness distribution, a stiff region larger than a single-cell size should trap the cells and tensional equilibrium can be realized (Scheme 2a). Besides, also in the case of a nano-patterned substrate with stiffness-heterogeneity, focal adhesions of the cells are preferentially formed on the dot where focal adhesions are more stabilized in a microscopic scale^44, 45, 46^, so the cells apparently sense such a substrate as an almost homogeneous field of stiffness, and finally reach tensional equilibrium^7, 47^. On the other hand, in the triangular patterns, the essential difference from these systems is that the cells actively remain away from tensional equilibrium in the process of nomadic migration among different regions of stiffness (Scheme 2b). The essential factor that makes this possible is the presence of dual competing taxis in the matrix. The system was arranged so that competing usual and reverse durotaxis can cause both the development and decrease of traction forces in an iterative manner. Therefore, fluctuations of the ISD can be amplified, which leads to the avoidance of tensional equilibrium.

**Scheme 2.**
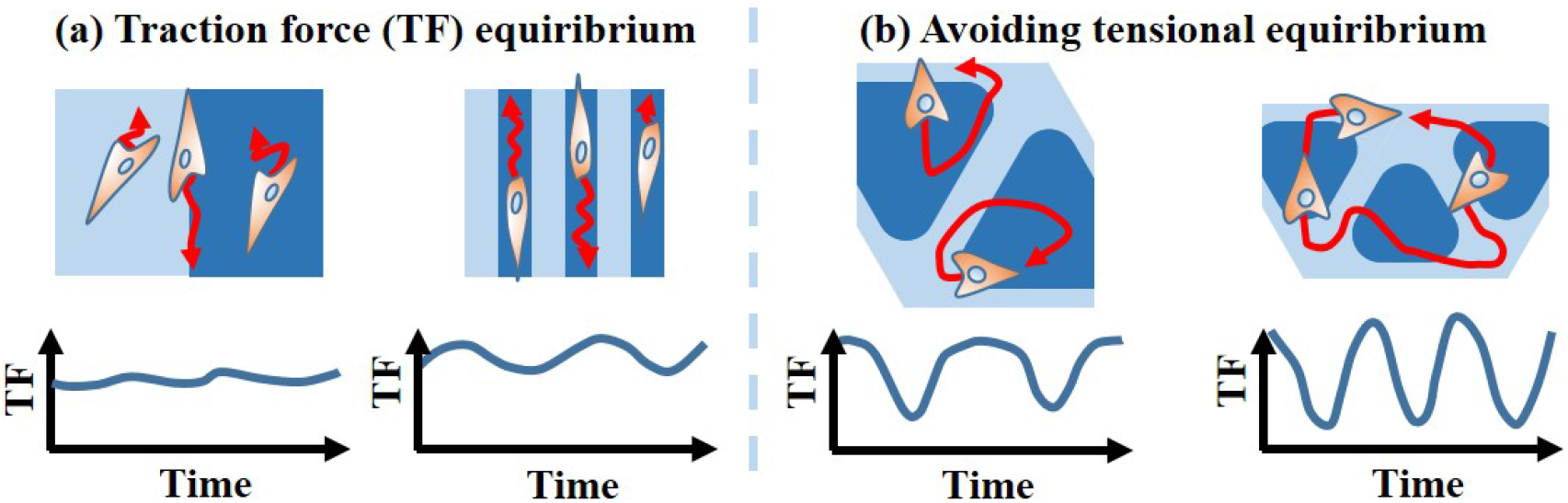
Schematic summary of traction force (TF) dynamics driven by the migration on the matrix with cell-scale stiffness-heterogeneity. **(a)** Cells on a striped patterned gel. Enhancement of TF fluctuation is limited because the cells are settling in a certain region. **(b)** Cells on a triangular patterned gel. TF fluctuation is significantly enhanced by nomadic migration.

Secondly, what kind of functional effects do migrating cells on a matrix with cell-scale stiffness-heterogeneity experience? The amplification of ISD fluctuations is linked to fluctuations in CSK tension and remodeling. This may have two major effects on cells. One effect is CSK-related mechanotransduction through signaling pathways including Rho-family GTPases^48^, Lats1/2 ^49, 50^, angimotin^51, 52^, and so on. Dynamic remodeling of CSK structures makes the intracellular localization of these factors strongly fluctuate, which causes frequent shuttling of nuclear translocation factors such as YAP/TAZ ^53, 54^ between the cytoplasm and nucleus. The other effect is the regulation of gene expression linked to nuclear deformation. It has been reported that intermingling of different chromosome territories mechanically occurs in response to cell shaping and nuclear deformation, and contributes to the regulation of gene expression^55^. The avoidance of tensional equilibrium should induce the shuttling of mechanosensitive nuclear translocating factors and the regulation of cellular function through ISD-dependent nuclear deformation. We will report complete set of our findings on functional modulation of the MSCs that experienced the far-from tensional-equilibrium and the related mechanical-stimuli-dependent regulation of gene expressions in another article soon.

Does nomadic cell migration on a heterogeneous mechanical field exist in a living organism? Some types of cells should migrate on micromechanically heterogeneous milieu, such as typically seen in wound healing of disordered tissues^56, 57^, in the biological development of immature organs^58, 59, 60^, and in stem cell localization inside a specific niche^61, 62^. In particular, MSCs are delocalized in niches composed of hard bone, soft vessels, and very soft blood cell surroundings in bone marrow, and are nomadically delocalized in the niche with such stiffness-heterogeneity^62^. This in vivo situation regarding nomadic cell migration, as inferred from the model system in the present study, should also induce the avoidance of tensional equilibrium.

## Conclusion

The present study revealed the principle conditions of cell-scale stiffness heterogeneity of ECM needed to enhance the fluctuations of ISD in cells migrating on it. The key factor for such enhancement is long-term continuous tactic cell movement, as exemplified in the coexistence of different cues of competing cellular taxis. Markedly enhanced ISD should lead to the avoidance of tensional equilibrium in an entire cell, which is expected to induce CSK-remodeling- and nuclear-deformation-linked functional regulation of migrating cells. This methodology should contribute to development of novel kind of sophisticated biomaterials which enable mechanobiology-based manipulation of cell functions.

## Author contributions

H. E. conducted all experiments, derivations, computations, and analyses. S. K. proposed the project, directed the research, and proposed the analyses. H. E. and S. K. discussed the results and wrote the paper.

## Data and materials availability

All data needed to evaluate the conclusions in the paper are present in the paper and/or the Supplementary Materials. Additional data related to this paper may be requested from the authors.

## Supplementary Information

### Rheological property of the StG gels

We investigated the rheological property of the StG gels. First, we measured the non-linear elasticity of the homogeneous StG gels by AFM (Fig. S1a). In these measurements, we used a silicone-nitride cantilever (PNP-TR, NanoWorld AG). We evaluated the dependence of the elasticity on the strain by changing the indentation depth of the gel with a cantilever of AFM under a constant maximum indentation force of 5 nN. We estimate the maximum stress *P_max_* through dividing the indentation force by contact area. Then, *P_max_* reached 3 – 30 kPa, which was comparable to the maximum traction stresses exerted by MSCs. The elastic modulus of the gels weakly depended on indentation depth and was almost constant. There were not observed clear shear thinning nor strain stiffening. This result indicated that the non-linearity of the elasticity of the StG gel was weak.

Next, we measured the dynamic viscoelasticity of the StG gels by AFM (Fig. S1b). The gel surface was indented with a cantilever with a trigger force 1 nN, then imposed with vertical oscillation of the constant amplitude of displacement (10 ~ 30 nm) and the frequency (1Hz). Through fitting the measured force amplitude and phase shift between displacement and force into the complex function derived from the Hertz model, we calculated the storage and loss modulus of the StG gels. For all the gels we used, the storage modulus was 10 – 20 times larger than the loss modulus, suggesting that StG gels are considered as the linear elastic body in the traction force microscopy.

**Fig. S1.**
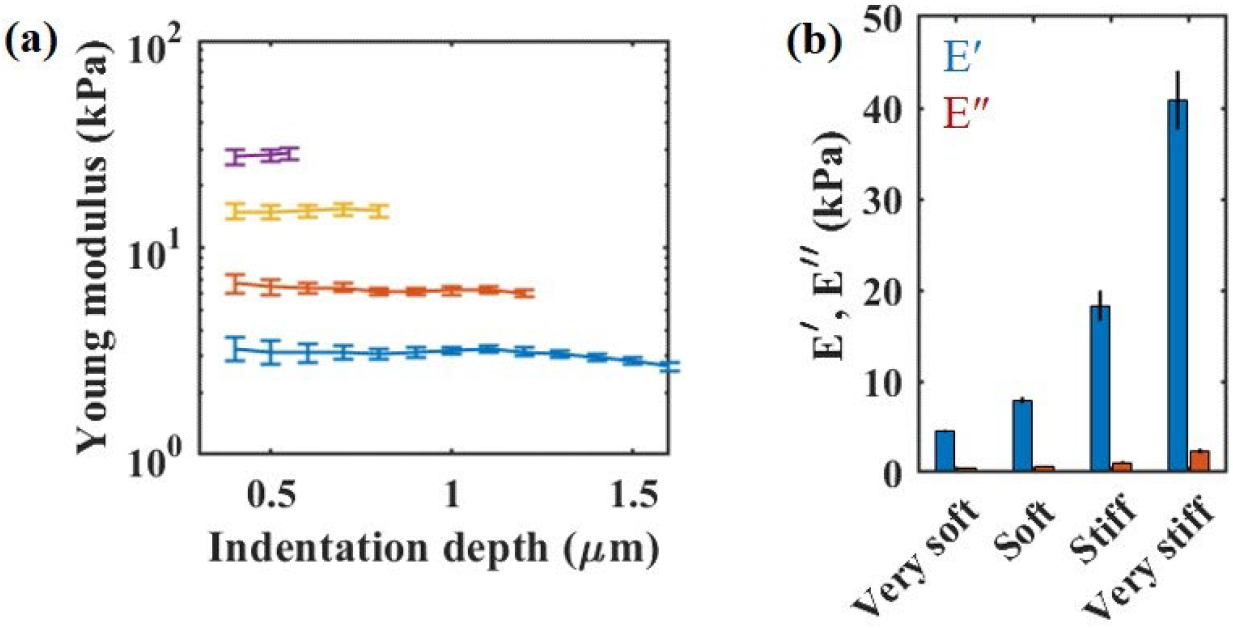
Rheological property of homogenous StG gels. (a) Dependence of the elasticity on the indentation depth of the StG gels. Colors indicate gels with different average elasticity. (b) Viscoelasticity of StG gels. Blue and red bars represent storage *E’* and loss *E”* modulus, respectively. Error bars denote standard deviation (n=5 for a and b).

### Topography of microelastically patterned gels

Topographies of the microelastically patterned gels were measured by AFM (Fig. S2). Height between bottom and top was typically 15 – 25 μm. Previously, we measured the durotactic behavior of MSCs on the gel with single and straight elasticity jump, which had similar topography^1^. In the case of a smaller elasticity jump than that of the threshold of durotaxis, we did not observe tactic motion from convex to concave regions even when the topography was around 25 μm. In addition, in the case of fibroblast on a striped gel with 100 μm unit width, cells freely migrate on the gel surface whose topography was around 10 μm, and no tactic motion was observed^2^. When we systematically changed the curvature of the elasticity boundary, fibroblast was not trapped in the concave regions^3^.

**Fig. S2.**
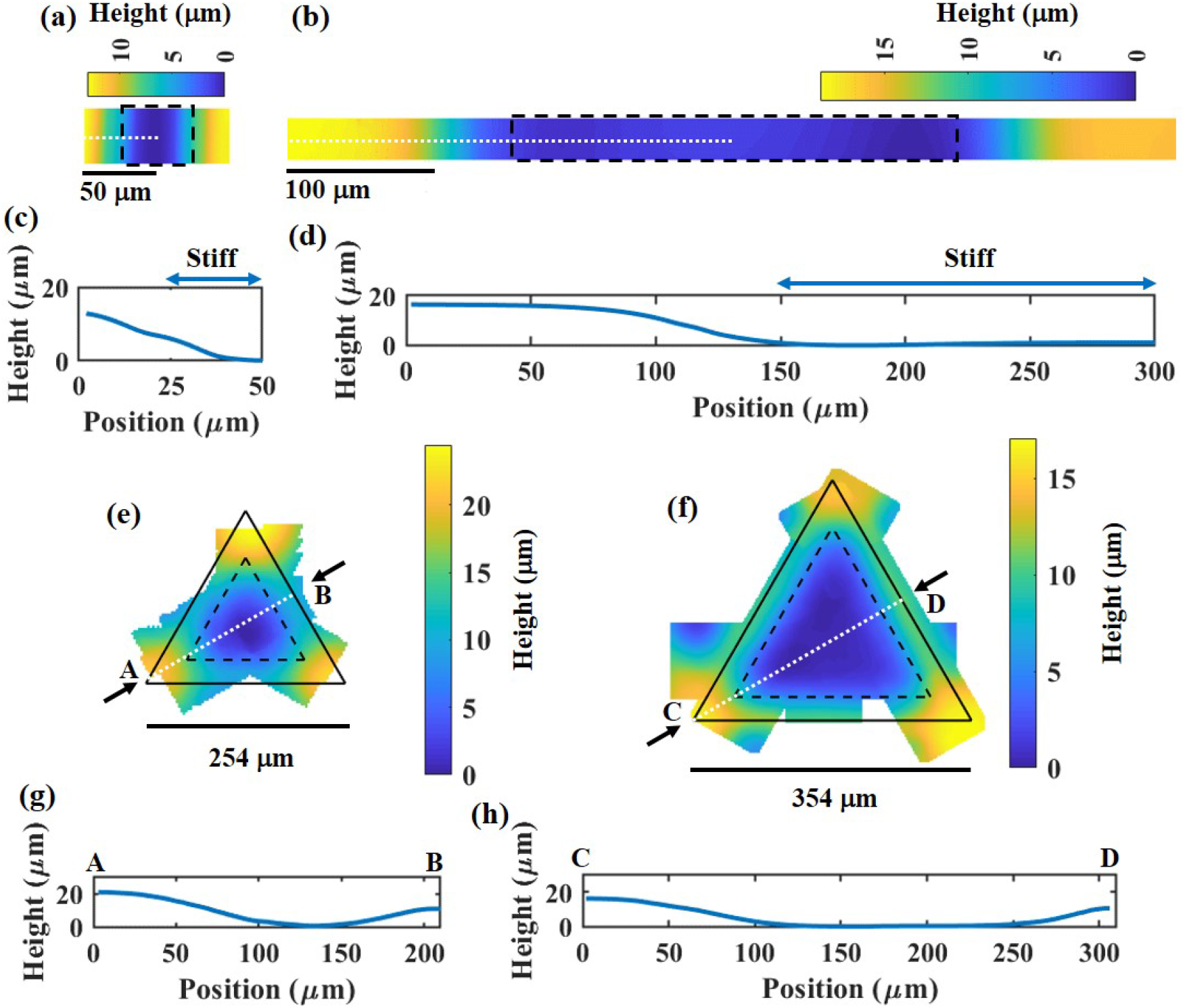
Topography of the microelastically patterned gels. Height distribution on stripe-100 (a, c), stripe-600 (b, d), triangle-150 (e, g), and triangle-250 gels (f, h). (a, b) Black dashed lines enclose the stiff area. (c, d) Cross section of the height distribution on stripe-100 (c) and stripe-600 (d). The height of the half period of the stripes is shown (white dashed line in a and b). (e, f) The solid line shows the unit triangle of the pattern. Dashed lines enclose the stiff area. (g, h) Cross section of the height distribution on Triangle-150 (c) and triangle-250 (d). White dashed lines in e and f indicate the position of the cross section. The arrows A – D in e and f represent the start and end points of the cross sections.

Topography was reported to affect the alignment of the cells. A previous study showed that the MSCs align in a longitudinal direction of the convex PDMS cylinder, while they randomly align on a concave cylinder^4^. In the case of striped StG gels, we found that the MSCs tended to align in the parallel direction against the stripes on both stiff stripes and soft stripes (Fig. S3). Especially, for a striped gel with 600 μm unit width, alignment became weaker and close to random on soft stripes. It was reported that the cells showed strong alignment against the stiff stripes^5^. Thus, the alignment of MSCs on striped StG gels could be driven by elasticity patterning. Model of durotactic cells showed that the strong alignment of the cells against the broad stiff stripe could cause the accumulation of the cells at the boundary between soft and stiff regions^2^, as described in Fig. 2a in the main text.

**Fig. S3.**
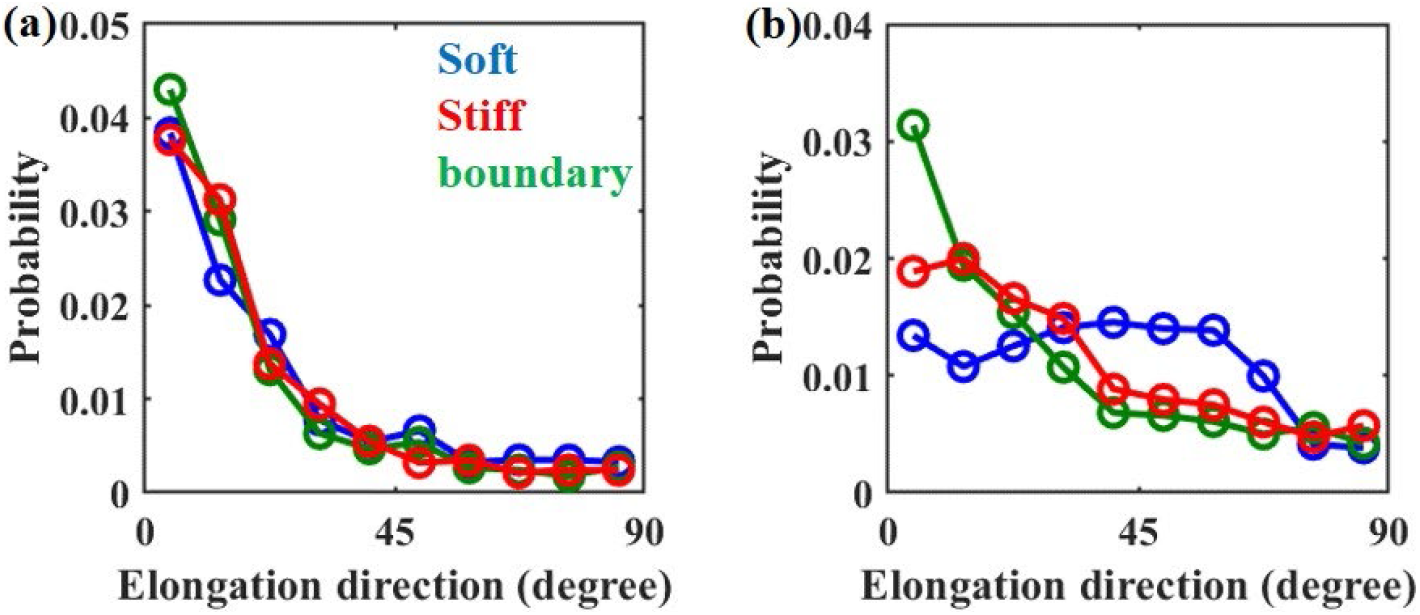
Alignment of MSCs on the striped pattern gels. The probability distribution of the direction of elongated cell body on stripe-100 (a) and stripe-600 gels. 0 degree indicates parallel direction against the stripes. We divided the stripe-patterned gels into soft (blue), boundary (green), and stiff (red) regions. (a) Young’s modulus of soft and stiff regions: 20 and 40 kPa, respectively. n = 66. (b) Young’s modulus of soft and stiff regions: 14 and 49 kPa, respectively. n = 93.

### Flow chart of FEM-based traction force microscopy on microelastically patterned gels

A flow chart of FEM-based traction force microscopy is shown in Fig. S4. The parameters and procedures that are not mentioned in the main text are as follows.

1. In the calculation of FEM, we used a hexahedron as a unit cell. The size of the hexahedron is 7.8 × 7.8 × 8.0 μm. The size of the imaginary gel used in the numerical calculation is 491.4 × 491.4 × 200 μm. The thickness of the microelastically patterned gel used in the experiment is around 200 μm.
2. To calculate the traction force from *M*(*x*, *x’*) and the measured displacement field, the origins of *M*(*x*, *x’*) and the displacement field should be matched. For this purpose, we calculated a height map of the fluorescent beads while making the full focus image, which reflected the topography of the surface. Because the topography and elasticity distribution had an almost inverse relation, we calculated the origin of the measured displacement field by pattern-matching between the inverse of height map and the distribution of elasticity used in the calculation of *M*(*x*, *x’*).
3. Since we only measured the displacement field of the gels in the *x* and *y* directions, we neglect the *z*-direction of the traction force and the topography of the gel surface as in usual 2-dimensional TFM.

**Fig. S4.**
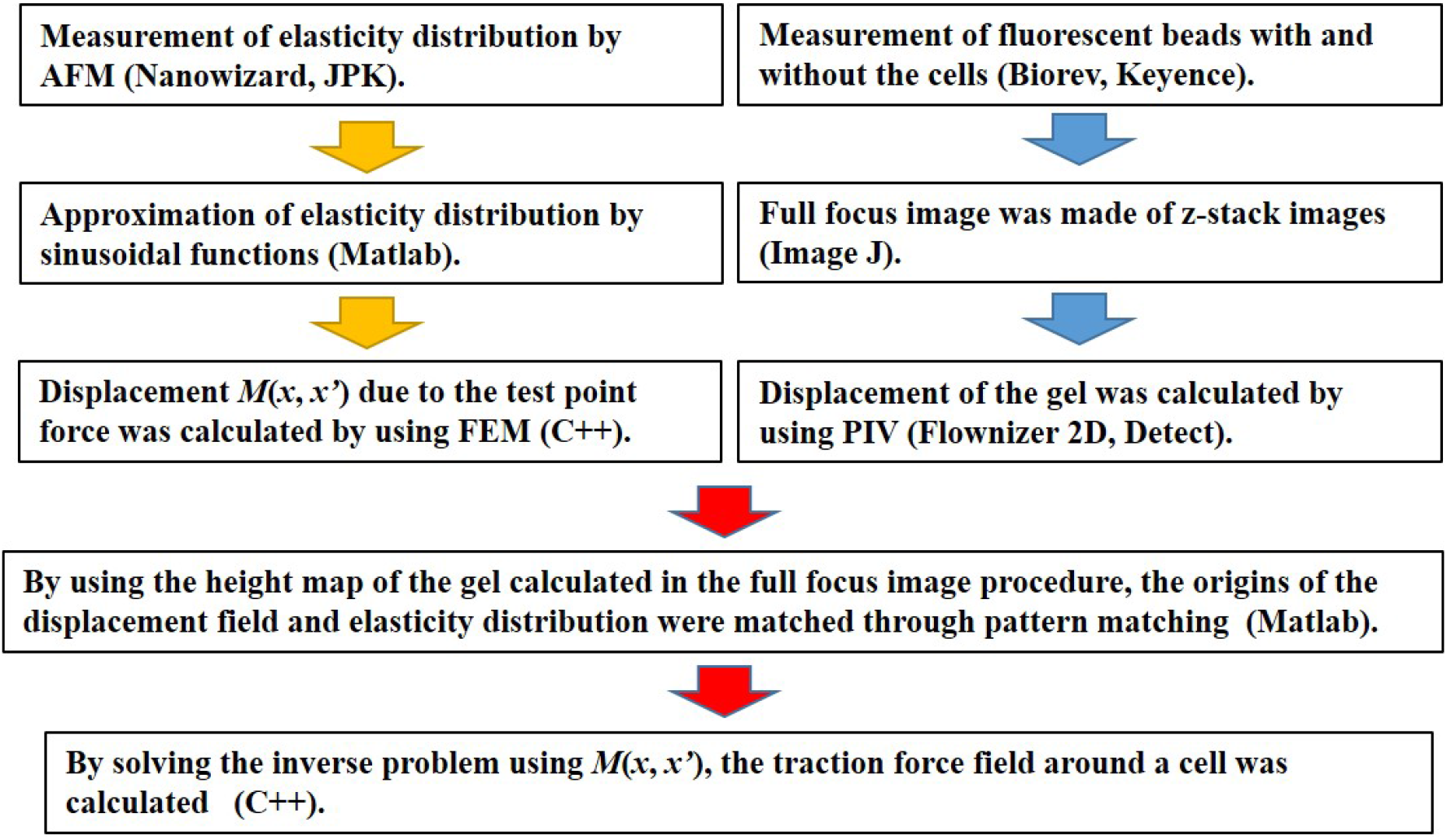
Flow chart of FEM-based traction force microscopy.

### Validity of FEM-based traction force microscopy

To check the validity of FEM-based traction force microscopy (FEM-TFM), we first compared the estimated traction force field from FEM-TFM and Fourier transform traction cytometry (FTTC) on homogeneous gels. FTTC is one of the most widely used methods for estimating the traction force. Figures S5 (a) and (b) show the estimated traction field by FEM-TFM (a) and the estimated traction field by FTTC (b). For FTTC, we adopted a Gaussian filter with a cut-off frequency of 0.066 μm^−1^ for the deformation field. The two methods give a very similar estimated traction field. However, for FEM-TFM, a large traction force tends to be localized in a narrow region. This phenomenon could be due to the regulation method we applied. Figure S5 (c) shows the time course of the mean traction stress under a cell. Blue and red lines represent the traction stress estimated by FEM-TFM and FTTC, respectively. The figure shows complete agreement between these two methods. For FTTC, the mean traction stress increases as the cut-off frequency of the Gaussian filter increases.

**Fig. S5.**
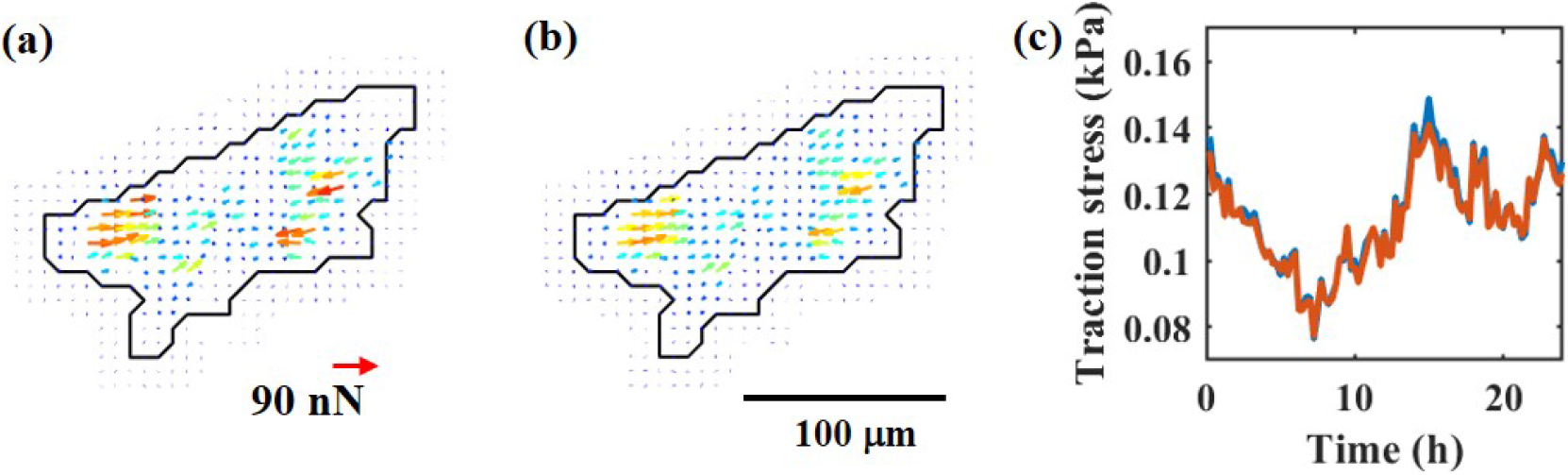
Traction force microscopy on homogeneous gels. (a) Field of the traction force reconstructed by FEM-TFM. (b) Field of the traction force reconstructed by FTTC. (a, b) Black line denotes the cell periphery. (c) Time course of mean traction stress under a cell. Blue: reconstruction by FEM-TFM. Red: reconstruction by FTTC. (a-c) Homogeneous soft gel. E = 5 kPa.

Next, through a numerical calculation, we checked the accuracy of the estimation method^6^ of the traction force on the elastically patterned substrate. First, we provided an artificial force distribution that mimicked traction forces exerted by a polarized MSC (Fig. S6 (a)). By using FEM, we calculated the deformation field at the surface of the substrate. Next, we added Gaussian noise to the deformation field with a standard deviation of 0.1 μm, which was the typical value of the background noise observed in the experiment (Fig. S6 (b)). Finally, we reconstructed the force distribution from the disturbed deformation field. As shown in Fig. S6 (c), we successfully estimated the force distribution. Compared to the forces in the soft region, estimated forces in the stiff region tended to be noisy, because the signal-to-noise ratio was lower in the stiff region.

**Fig. S6.**
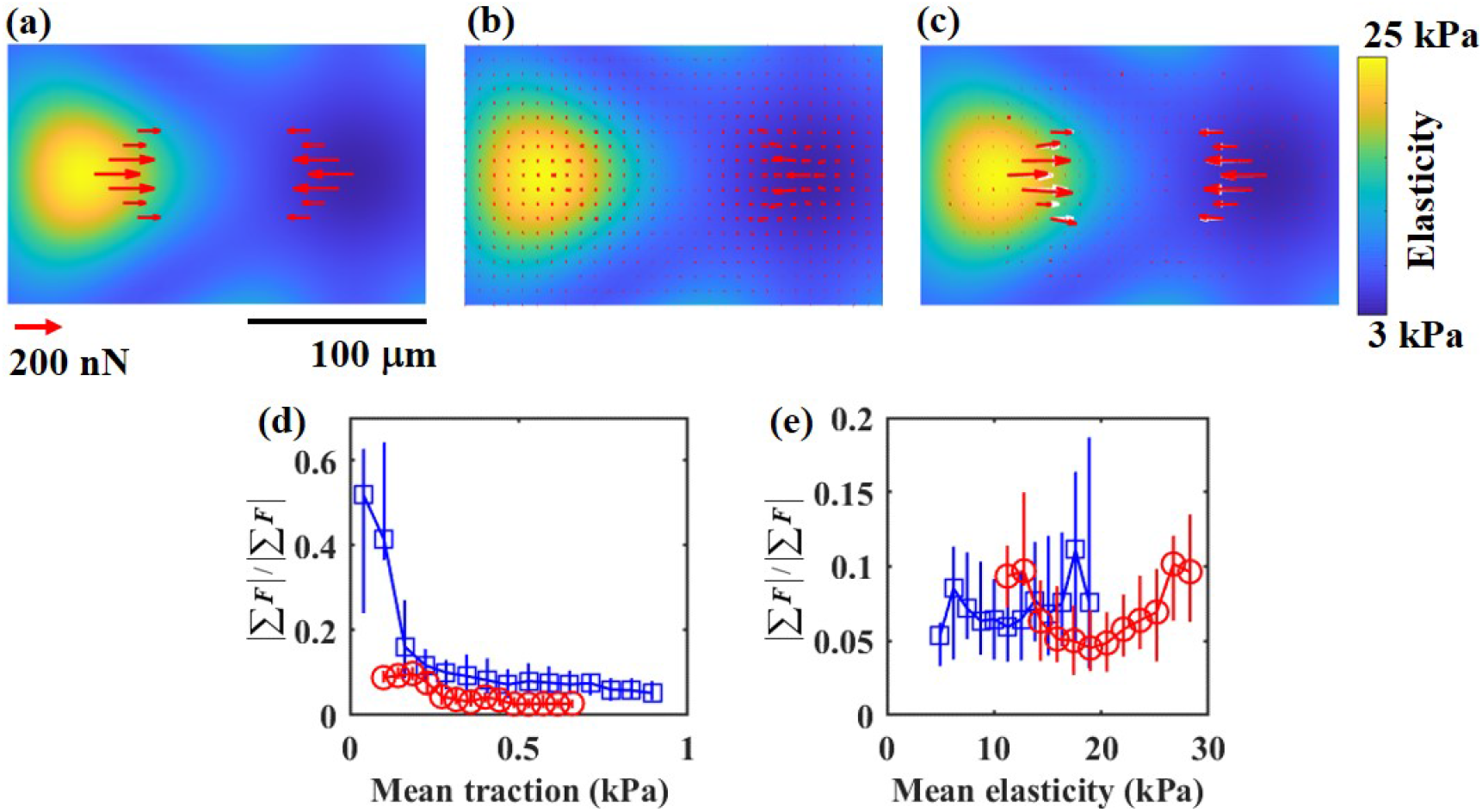
Traction force microscopy on microelastically patterned gels. (a – c) Test of force reconstruction by numerical calculation. (a) Test forces that mimic the forces from the polarized MSCs. (b) Calculated deformation field at the surface of the gel by using the finite element method. We added white Gaussian noise to the deformation field, where the standard deviation of the noise is 0.1 μm. (c) Estimated force distribution (red arrow) from the disturbed deformation field, Fig. 3 (b). White arrow represents the true forces. (d, e) Dependence of the magnitude of the unbalance of the forces on the mean traction stress of a cell (d) and the mean elasticity under a cell (e). (d) Blue: Triangular-150. Red: Plain-stiff. (e) Blue: Triangle-150. Red: Stripe-100.

Finally, we briefly examined the accuracy of FEM-TFM for cells on the microelastically patterned gels. It has been reported that the unbalance of the traction force vectors |∑ *F*| / ∑| *F*| became nearly zero on homogeneous gels^7, 8^, because the inertial force and viscous drag force are almost negligible for adhesive cells. Thus, a condition of zero-sum force should also be satisfied on elastically patterned gels. In general, |∑ *F*| / ∑| *F*| has a finite value due to the error of the measurement of the displacement field and the procedure used for force reconstruction^7, 8^. Thus, the magnitude of unbalance of the forces, |∑ *F*| / ∑| *F*|, is a measure of the accuracy of TFM. For all experimental data, average |∑ *F*| / ∑| *F*| are around 2.9 % and 5.7 % for homogeneous and patterned gels, respectively. As shown in Fig. S6 (d), for both homogeneous and patterned gels, unbalance of the forces is negatively correlated with the magnitude of the mean traction stress (correlation coefficient ~ −0.4 – −0.3). Especially, when the traction force is too small (~0.1 kPa), the force distribution was hardly estimated. On the other hand, the mean elasticity under the cell had almost no correlation with the unbalance of the forces (correlation coefficient ~ 0 – 0.1, Fig. S6 (e)). Thus, in our experiment, the accuracy of force reconstruction could not significantly depend on the spatial variation of elasticity.

### Threshold of the force spot calculated from the cumulative distribution

To identify the force spot, we calculated the threshold of the traction stress. To define this threshold, we used the complementary cumulative distribution (CFD, *C*(*τ*)) of the traction stress. The derivative of CFD provides the probability distribution function. As shown in Fig. S7, CFD sharply decreased at a small traction stress, which comes from the background noise. CFD had exponential tails, as reported in the previous work on a homogeneous gel. Thus,

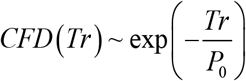

is satisfied for large traction stress. The exponential tails should be due to the traction force exerted by a cell. Thus, we use typical stress *P_0_* as the threshold for the force spot.

**Fig. S7.**
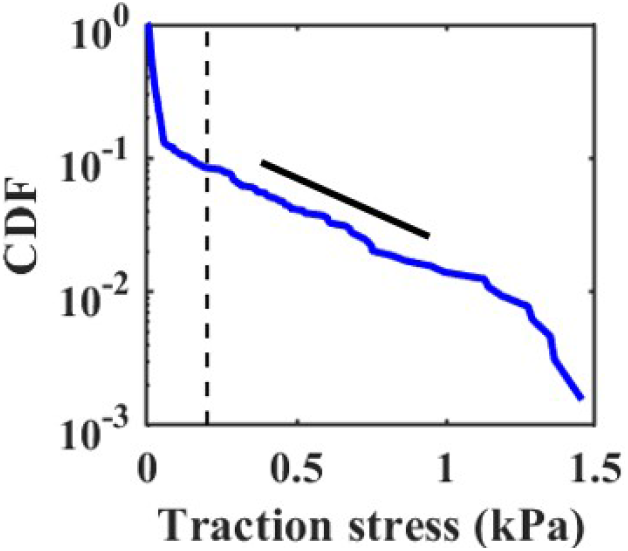
Complementary cumulative distribution of the traction stress of a cell at a certain time.

### Magnitude of traction stress in force spots on homogeneous gels

Figures S8 (a) and (b) show the time series of the field of traction force on Plain-soft and Plain-stiff. Cells on Plain-soft had a rather circular shape, while cells on Plain-stiff had an elongated shape. As shown in Fig. S8 (c), traction stress in the force spot on Plain-stiff was significantly greater than that on Plain-soft.

**Fig. S8.**
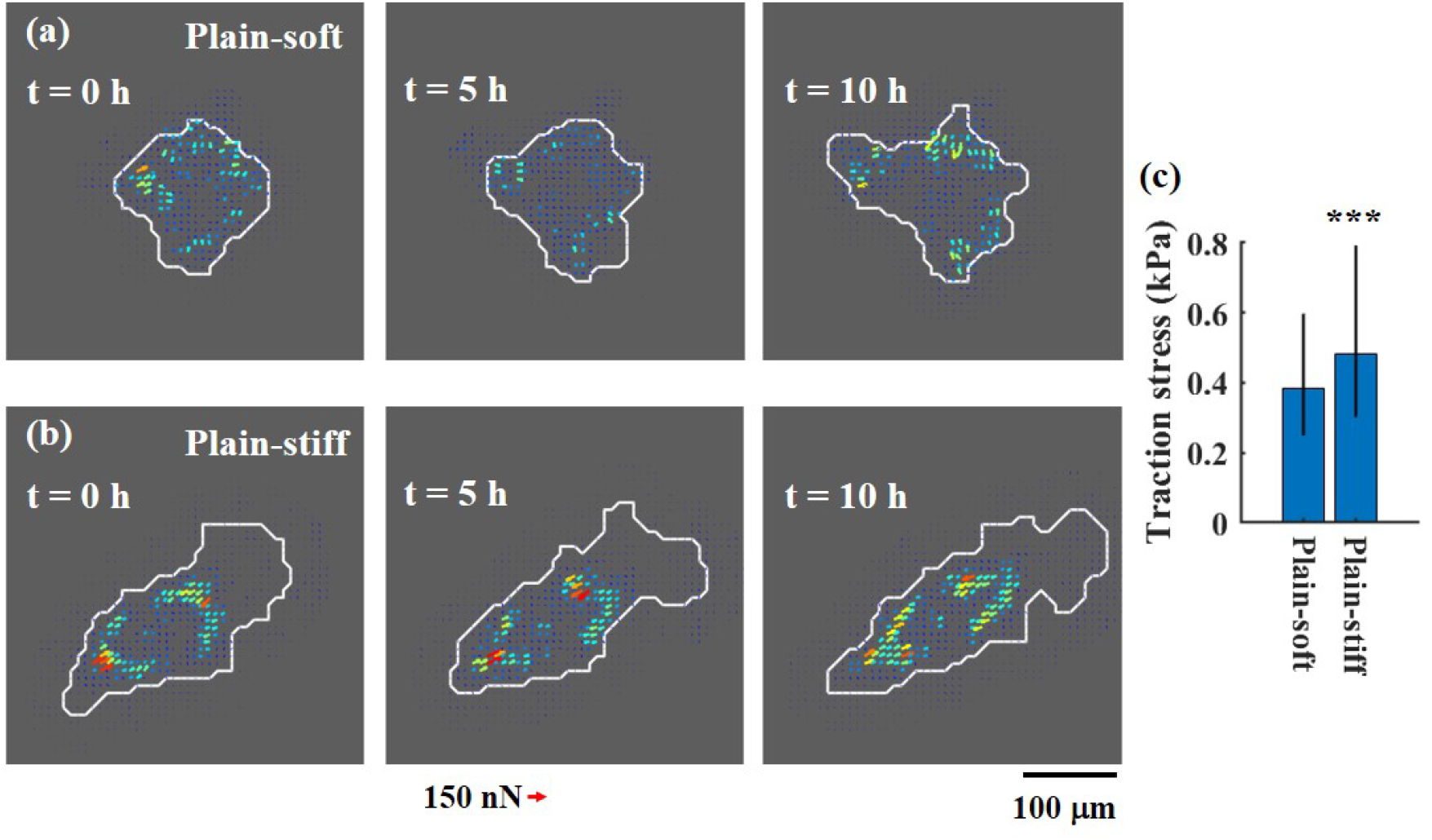
Traction forces on the control homogeneous gels. (a, b) Time series of the traction force field on Plain-soft (a) and Plain-stiff (b). (c) Bar graph of the magnitude of traction stress in force spots. Bar denotes the median value. Error bar connects the 0.25 and 0.75 quantiles. The Mann–Whitney U test was used to calculate the P value. *** P < 0.001. Plain-soft: n = 39. Plain-stiff: n=28.

### Residence time in soft and stiff regions

We calculated the residence time, which is the amount of time a cell centroid remained in the soft and stiff regions (Fig. S9). Here, we defined the irradiated region of the photomask as a stiff region and the masked region as a soft region (Fig. 1 (c) and (d) inset). Residence times in soft and stiff regions were almost equal for wide stripes and a small triangular pattern. Otherwise, MSCs stayed in stiff regions longer. Compared to the residence time on stripe patterns, the residence time on triangular patterns was short, which indicates that the cell frequently moved between soft and stiff regions.

**Fig. S9.**
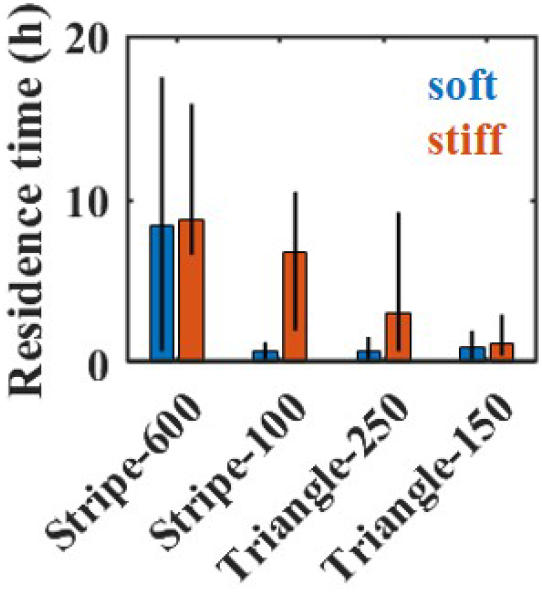
Residence time of cells in the soft and stiff regions on each patterned gel. Bar denotes the median value. Error bar connects the 0.25 and 0.75 quantiles. Stripe-600: striped patterned gel with L = 600 μm. Stripe-100: striped patterned gel with L = 100 μm. Triangle-250: triangular patterned gel with L = 250 μm. Triangle-150: triangular patterned gel with L = 150 μm. Stripe-600: n = 25. Stripe-100: n = 32. Triangle-250: n = 23. Triangle-150: n = 46.

### Relation between the fluctuations of mean traction stress and of mean elasticity

We examined whether the feature of fluctuation of traction stress during nomadic cell migration among different regions of stiffness was able to be persisted across the elasticity boundary or not, i.e., memory effect on the stress fluctuation persisted from the previous region. To answer this issue, we investigated the detailed relation between the fluctuation of traction stress and of mean elasticity under the cell with a time scale of a few hours. Here, we divided the time courses of mean elasticity that the cell senses and of generated traction stress into the time window of 4 hours (Fig. 4a-c in the main text). Then, we calculated fluctuation of traction stress *Fluc*_4*h*_ (*Tr*) = σ_4*h*_ (*Tr_m_*) / ⟨*Tr_m_*⟩ and fluctuation of the mean elasticity under the cell *Fluc*_4*h*_ (*E*) = σ_4*h*_ (*E_m_*) / ⟨*E_m_*⟩, where σ_4*h*_ (•) and ⟨•⟩ represent standard deviation and time average of variables in the time window of 4 hours (see Method). *Fluc*_4*h*_ (*E*) = 0 indicates that the mean elasticity under the cell is constant, and the cell migrates around the region with the same elasticity. On the homogeneous gels, *Fluc*_4*h*_ (*E*) = 0 is always satisfied. On the other hand, *Fluc*_4*h*_ (*E*) ≫ 0 gives the situation that the cell shows transboundary motion between soft and stiff regions.

To clarify how the short-time fluctuation of mean elasticity affected the fluctuation of traction stress, we measured the correlation between *Fluc*_4*h*_ (*E*) and *Fluc*_4*h*_ (*Tr*). As shown in Fig. S10a, we found that *Fluc*_4*h*_ (*Tr*) had a positive correlation with *Fluc*_4*h*_ (*E*) (correlation coefficients 0.25 – 0.45). To clarify the detailed relation, we showed averaged values of *Fluc*_4*h*_ (*Tr*) in the sections, 0.02(*n* − 1) < *Fluc*_4*h*_ (*E*) < 0.02*n*, *n* = 1,2,3… (Circles and crosses in Fig. S10a). The averaged values indicated linear relation between the fluctuations. The linear positive relation meant that the fluctuation of traction stress became larger when the cell migrated between soft and stiff regions. The linear fit between *Fluc*_4*h*_ (*Tr*) and *Fluc*_4*h*_ (*E*) gave how the fluctuation of traction stress depended on the time variation of mean elasticity (Black dashed lines in Fig. S10a). The slope of the line indicates how much sensitively fluctuation of traction stress responds to that of mean elasticity, and the intercept of *Fluc*_4*h*_ (*Tr*) axis gives the estimation of the degree of fluctuation of the traction stress with constant mean elasticity.

To investigate the persistence of the stress fluctuation driven by the nomadic migration, we focused on the intercept of linear fit between *Fluc*_4*h*_ (*Tr*) and *Fluc*_4*h*_ (*E*). When the large stress fluctuation persisted, the intercept should be larger than the degree of stress fluctuation on homogeneous gels. In Figs. S10b and c, we showed the median value of the fluctuation of the traction stress on homogeneous gels (Fig. S10 b), and the intercepts of linear fit between *Fluc*_4*h*_ (*Tr*) and *Fluc*_4*h*_ (*E*) on the patterned gels (Fig. S10 c). The intercepts of striped patterned gels had close value to *Fluc*_4*h*_ (*Tr*) on homogeneous gels, while the intercepts of triangular patterned gels had much larger values. This result meant that the fluctuation of traction stress on striped patterned gels behaved similarly to that on the homogeneous gels, when the cell incidentally migrated on the region with the same elasticity. On the other hand, the fluctuation of traction stress on triangular patterned gels kept high value for a few hours, even when the cells did not temporarily move between soft and stiff regions (Fig. S10c). Thus, it suggested that the fluctuation of traction stress on triangular patterned gels kept the memory of the time variation of the mean elasticity for a few hours, while that on the striped patterned gels immediately loss the memory.

**Fig. S10.**
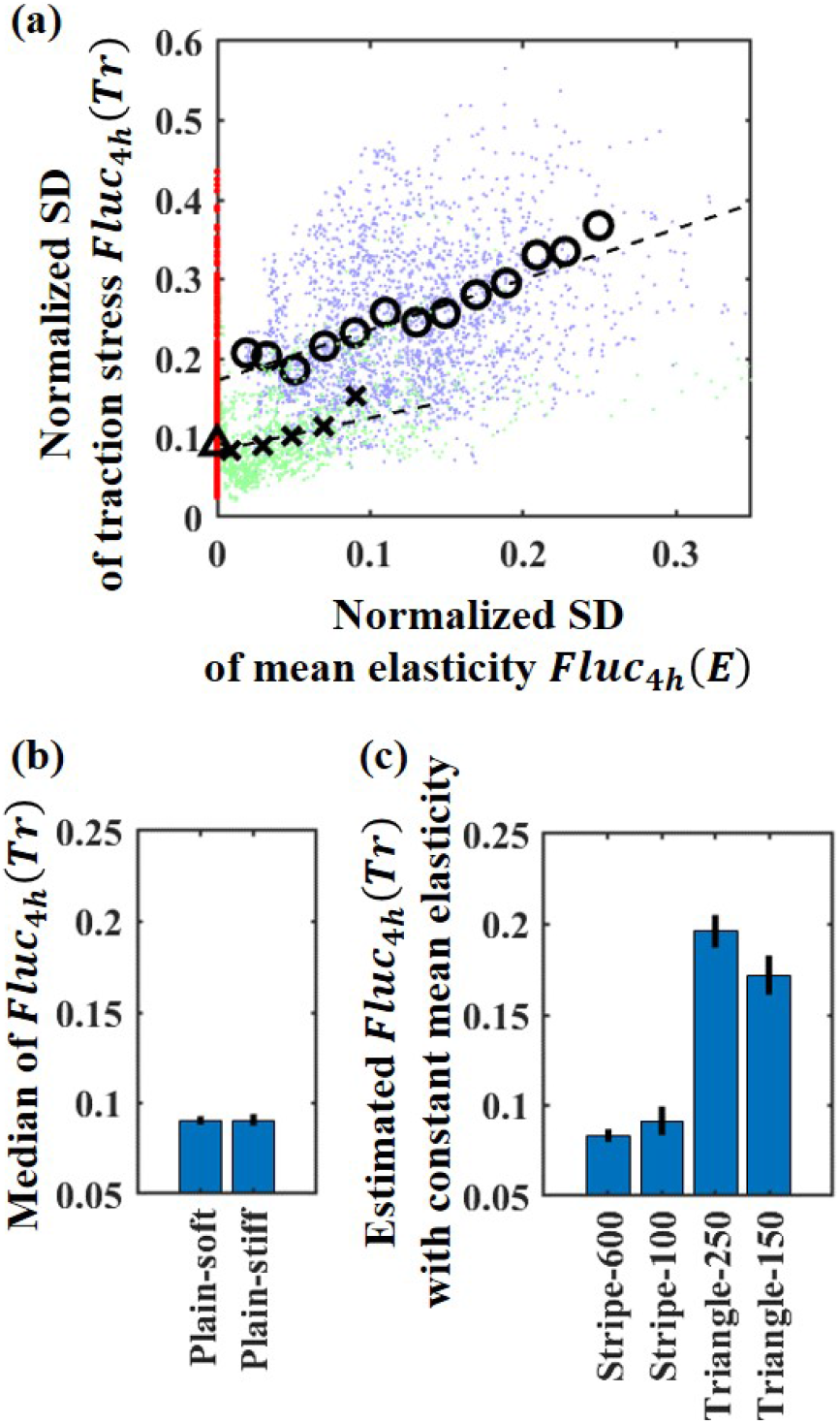
Relation between the fluctuations of mean traction stress and of mean elasticity. **(a)** Correlation between *Fluc*_4*h*_ (*E*) and *Fluc*_4*h*_ (*Tr*). Red dot: Plain-stiff. Green dot: Stripe-600. Blue dot: Triangle-150. Black dashed line: linear fit of the data. Triangular symbol: median of Plain-stiff. Circles and crosses: mean values in the section, 0.02(*n* – 1) < *Fluc*_4*h*_ (*Tr*) < 0.02*n*, *n* = 1,2,… **(b)** Median of *Fluc*_4*h*_ (*Tr*) on homogeneous gels. **(c)** Estimated degree of fluctuation of the traction stress with constant mean elasticity. The intercepts of the linear fit between *Fluc*_4*h*_ (*E*) and *Fluc*_4*h*_ (*Tr*) are shown. **(b, c)** Error bars represent 95% confidential intervals. Plain-soft: n = 39. Plain-stiff: n=28. Stripe-600: n = 25. Stripe-100: n = 32. Triangle-250: n = 23. Triangle-150: n = 46.

